# Comprehensively characterizing evolutionarily conserved differential dosage compensations of the X chromosome during development and in diseases

**DOI:** 10.1101/2022.07.25.501478

**Authors:** Mengbiao Guo, Zhengwen Fang, Bohong Chen, Zhou Songyang, Yuanyan Xiong

## Abstract

The active X chromosome in mammals is upregulated to balance its dosage to autosomes during evolution. However, it is elusive why the currently known dosage compensation machinery showed uneven and small influence on X genes, necessitating systemic investigation of X dosage in different angles and identification of new dosage regulators. Here, based on >20,000 transcriptomes, we identified two X gene groups (genome ploidy-sensitive [PSX] and ploidy-insensitive [PIX] genes), showing distinct but evolutionarily-conserved (in both primates and mouse) dosage compensations (termed X-over-Autosome dosage Ratio, or XAR). We then explored XAR in diseases and in stem cells, where XAR is potentially important. We demonstrated that XAR-PIX was downregulated while XAR-PSX upregulated across cancer types at both RNA and protein levels. In contrast, XAR-PIX was upregulated while XAR-PSX downregulated during stem cell differentiation. Interestingly, XAR-PIX, but not XAR-PSX, was significantly lower and associated with autoantibodies and inflammation in lupus patients, suggesting that insufficient dosage of PIX genes may contribute to lupus pathogenesis. We further identified and experimentally validated two new XAR regulators, *TP53* and *ATRX*. Collectively, we provided insights to further unravel the mystery of X dosage compensation in mammals and its pathophysiological roles in human diseases.

## Introduction

In mammals, females have two X chromosomes while males have only one and females evolved to silence one copy of X to match the single X copy in males, a phenomenon called X chromosome inactivation (XCI) (Disteche 2012) with exceptions of escaping genes (Dunford, et al. 2017). Initiated early in female embryo development, XCI is essentially stable for life after complete (Vallot, et al. 2016). To balance the dosage between the single active X chromosome and other autosomes with two active copies, both sexes evolved to upregulate the active X chromosome (Disteche 2012; Deng, et al. 2013; Larsson, et al. 2019), although with controversies (Xiong, et al. 2010; Julien, et al. 2012).

Meanwhile, some possible mechanisms underlying X upregulation in mammals were revealed, mainly through epigenetic regulation, including transcription elongation and initiation enhancement and RNA stability improvement (Deng, et al. 2013). Expectedly, *KAT8* (lysine acetyltransferase 8, also known as *MOF* or *MYST1*), *MSL1* (male-specific lethal 1 homolog), and *MSL2*, whose homologs are responsible for X upregulation in Drosophila, were found to behave similarly in mammals and regulate histone acetylation to enhance transcription preferably on X chromosome (Deng, et al. 2013). However, it is intriguing that *KAT8* affected X genes unevenly and only a small percentage of X genes showed considerable effect sizes.

Recently, using published single-cell RNA sequencing datasets, Larsson and colleagues reported that the frequency of transcriptional bursts was the driving force behind X upregulation in mouse, and they proposed that trans-acting enhancer-binding regulators may be the underlying factor behind the higher transcriptional burst frequency of X chromosome (Larsson, et al. 2019). However, these factors were unknown and the hypothesis of enhancer factors in X upregulation was not verified.

*TP53* is a highly-conserved gene important for various stress responses (Lu, et al. 2009). It was reported to interact with key MSL components, *KAT8* and *MSL2* (Kruse and Gu 2009; Li, et al. 2009). The p53 binding sites (Nguyen, et al. 2018) in *KAT8* (one site in promoter) and *MSL2* (three sites in promoter) indicate direct regulation of *KAT8* and *MSL2* transcription by p53. ATRX, an X-linked telomere-associating protein studied by our team (He, et al. 2015; Hu, et al. 2016), is a chromatin remodeler that functions in genome stabilization and heterochromatin formation (Dyer, et al. 2017; Teng, et al. 2021). ATRX works with XIST and PRC2 to remodel chromatins (Ren, et al. 2020) and ATRX loss can affect X inactivation in embryonic trophoblast (Garrick, et al. 2006). Importantly, it has been reported that TP53 and ATRX (also known as RAD54) physically interact with each other (Linke, et al. 2003) and work together in the same pathway (Seah, et al. 2008; Oppel, et al. 2019; Gulve, et al. 2022). Both genes are involved in genome-wide histone modifications (Allison and Milner 2004; Dyer, et al. 2017), thus capable of regulating a large number of genes. We therefore hypothesized that TP53 and ATRX probably regulate X dosage compensation.

Interestingly, *TP53* mutations contributed to the sex discrepancy of cancer incidences by differentially regulating X-linked genes between two sexes (Haupt, et al. 2019). Notably, *TP53* mutations are the most frequent events across all cancers (Hainaut and Pfeifer 2016), and more than 91% of tumors with *TP53* mutations show functional loss of both *TP53* copies (Donehower, et al. 2019). p53 is also implicated in immune regulation of autoimmune diseases (Munoz-Fontela, et al. 2016). On the other hand, X-linked genes have been implicated in many diseases, including cancer (Haupt, et al. 2019) and autoimmune diseases like systemic lupus erythematosus (SLE) (Hewagama, et al. 2013). Abnormal X dosage has also been reported in both cancer and SLE (Pageau, et al. 2007; Syrett, et al. 2019). For example, X upregulation has been contributed to the loss of Barr body (the inactivated X chromosome) in breast and ovarian cancers. X dosage also plays an important role in cancer initiation, because excessive X upregulation by somatic X reactivation under chronic stress leads to cancer (Yang, et al. 2020). However, the dynamics of X-over-autosome dosage ratios (XAR) remain to be studied systematically across cancers, which accumulates a large number of high-throughput sequencing datasets, for example, The Cancer Genome Atlas (TCGA) (Davoli, et al. 2013). More importantly, large numbers of cancer mutations and genome-wide expression profiles from TCGA have the potential to reveal new regulators of XAR, including both protein-coding and long noncoding RNA (lncRNA) genes, to uncover the elusive mechanism of dosage compensation in humans. However, these possibilities have not been explored yet.

In this work, we identified two distinct groups of genes (PIX and PSX) on X chromosome and characterized their dosage compensations in about 20,000 RNA sequencing (RNA-Seq) samples from TCGA and the Genotype-Tissue Expression (GTEx) project (GTEx-Consortium 2013), and thousands of proteomic samples from Clinical Proteomic Tumor Analysis Consortium (CPTAC). We provided a systemic view of both XAR-PIX and XAR-PSX in human. We showed that the differential dosage between PIX and PSX genes were evolutionarily conserved. Interestingly, XAR-PIX and XAR-PSX were affected differently in sex-biased diseases and during stem cell differentiation which is closely linked with XCI (Hall, et al. 2008). Lastly, we identified and experimentally validated two new XAR regulators, *TP53* and *ATRX*. In summary, our work contributed to better understanding of X dosage compensation under different conditions.

## Results

### Two groups of X chromosome genes with distinct sensitivities to genome ploidy

We are interested in which genes on X chromosome require less dosage compensation, since only a subset of X chromosome genes were affected after *KAT8* knockdown (Deng, et al. 2013). To this end, we examined X chromosome gene sensitivities to genome ploidy, defined by the absolute Pearson’s correlation coefficients between protein-coding gene expression levels (n=615 X chromosome genes and n=16,622 autosome genes remained, **Table S1**) with genome ploidies of cancer samples from TCGA. Based on the approximately normal distribution of these correlations, we identified two groups of X chromosome genes, termed ploidy-sensitive X chromosome (PSX) genes, and ploidy-insensitive X chromosome (PIX) genes (see **Methods**), separated by the mean of the correlation distribution (**Fig. 1A**). As expected, PSX and PIX genes showed distinct expression distribution. PSX genes (n=286, **Table S1**) had highly similar expression pattern to autosome genes (**Fig. 1B**), while PIX genes (n=329, **Table S1**) had much lower expression levels, with a median of about 2/3 of PSX genes expression (**Fig. 1C**). Of note, no preference for genes escaping X inactivation was observed in PSX or PIX. Interestingly, we found three PIX-specific gene clusters on X chromosome, compared to PSX genes (**Fig. 1D**). The PIX cluster located at around 100Mb (hg38) contained eight out of nine Transcription Elongation Factor A Protein-Like (TCEAL) genes, encoding nuclear phosphoproteins.

**Fig. 1.**
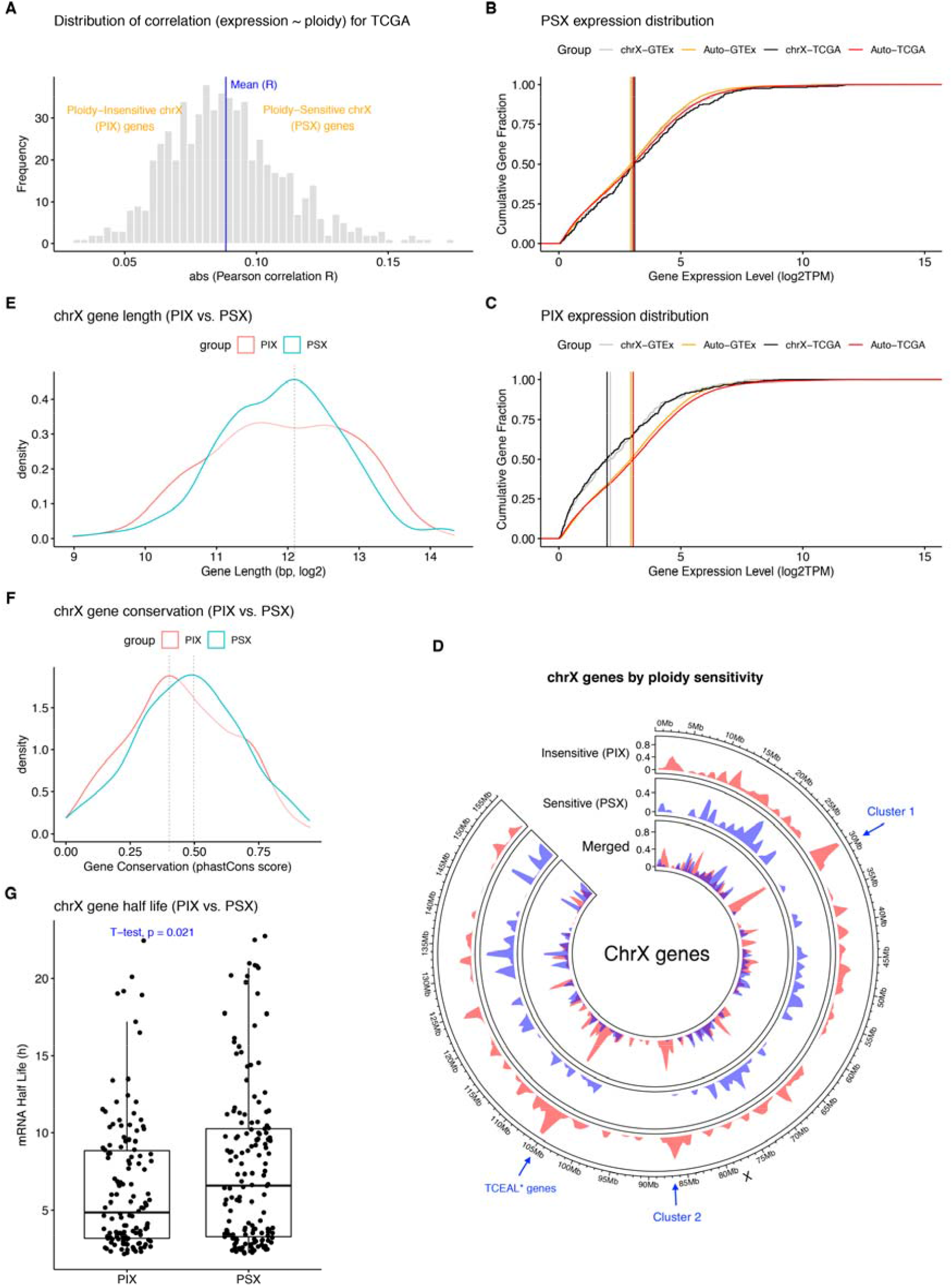
Identification and characterization of PSX and PIX genes on X chromosome. (**A**) Classification X chromosome genes based on absolute values of Pearson correlations (x-axis) compared to the average absolute correlation value (blue vertical line). (**B-C**) Cumulative distribution of expression levels of PSX (**B**) and PIX (**C**) genes on X chromosome, compared to autosomal genes, using samples from the TCGA project or the GTEx project. A total of 100 samples were randomly selected and their median expression levels for genes were shown for visualization. Vertical lines indicate medians. (**D**) PIX and PSX genes distribution on X chromosome. PIX enriched regions were marked by arrows. (**E-G**) Comparing gene length (**E**), gene sequence conservation (**F**), and mRNA half-life (two-sided *t*-test, **G**) between PIX and PSX genes. PIX: ploidy-insensitive X chromosome genes, PSX: ploidy-sensitive X chromosome genes.

### Different features and regulation of PIX and PSX genes

The distinct expression pattern between PIX and PSX genes prompted us to explore their differential genomic features. First, we found that the lengths (excluding introns) of PIX genes spanned a large range from 2kb to 8kb, while lengths of PSX genes peaked at around 4kb (**Fig. 1E**). This trend was similar when including introns (**Fig. S1A**). Although GC content of mRNAs is critical for their storage and decay (Courel, et al. 2019), we found similar GC content between PIX and PSX genes (**Fig. S1B**). Moreover, we observed lower conservation of PIX genes than PSX genes (**Fig. 1F**), which resembled the faster-X effect in non-coding regions on X chromosome (Meisel and Connallon 2013). RNA half-lives were reported to be longer for X-linked than autosomal genes (Deng, et al. 2013). Inspired by this, we examined the mRNA half-lives of PIX and PSX genes and found that PIX (4.9h) tended to have shorter mRNA half-lives than PSX genes (6.6h) (**Fig. 1G**), consistent with lower expression levels of PIX than PSX genes.

### Evolutionally conserved differential dosage compensation between PIX and PSX genes

Based on these two groups of X chromosome genes (PIX and PSX), we calculated two dosage ratios of X chromosome over autosome (XAR, see **Methods**) to evaluate their dosage compensation, termed XAR-PIX and XAR-PSX, respectively. As expected, XAR-PSX dosages were around one across GTEx tissues (**Fig. 2A**), but mostly above one across TCGA cancer types (**Fig. 2B**). In contrast, XAR-PIX dosages were much lower than XAR-PSX and fluctuated around 0.6 both across GTEx tissues and across TCGA cancer types. Furthermore, the difference between XAR-PIX and XAR-PSX was also observed in non-human primates (**Fig. S2A**) and mouse (**Fig. S2B**) across tissues, except for the brain. It suggests that PIX and PSX gene classification and their dosage compensation are evolutionally conserved.

**Fig. 2.**
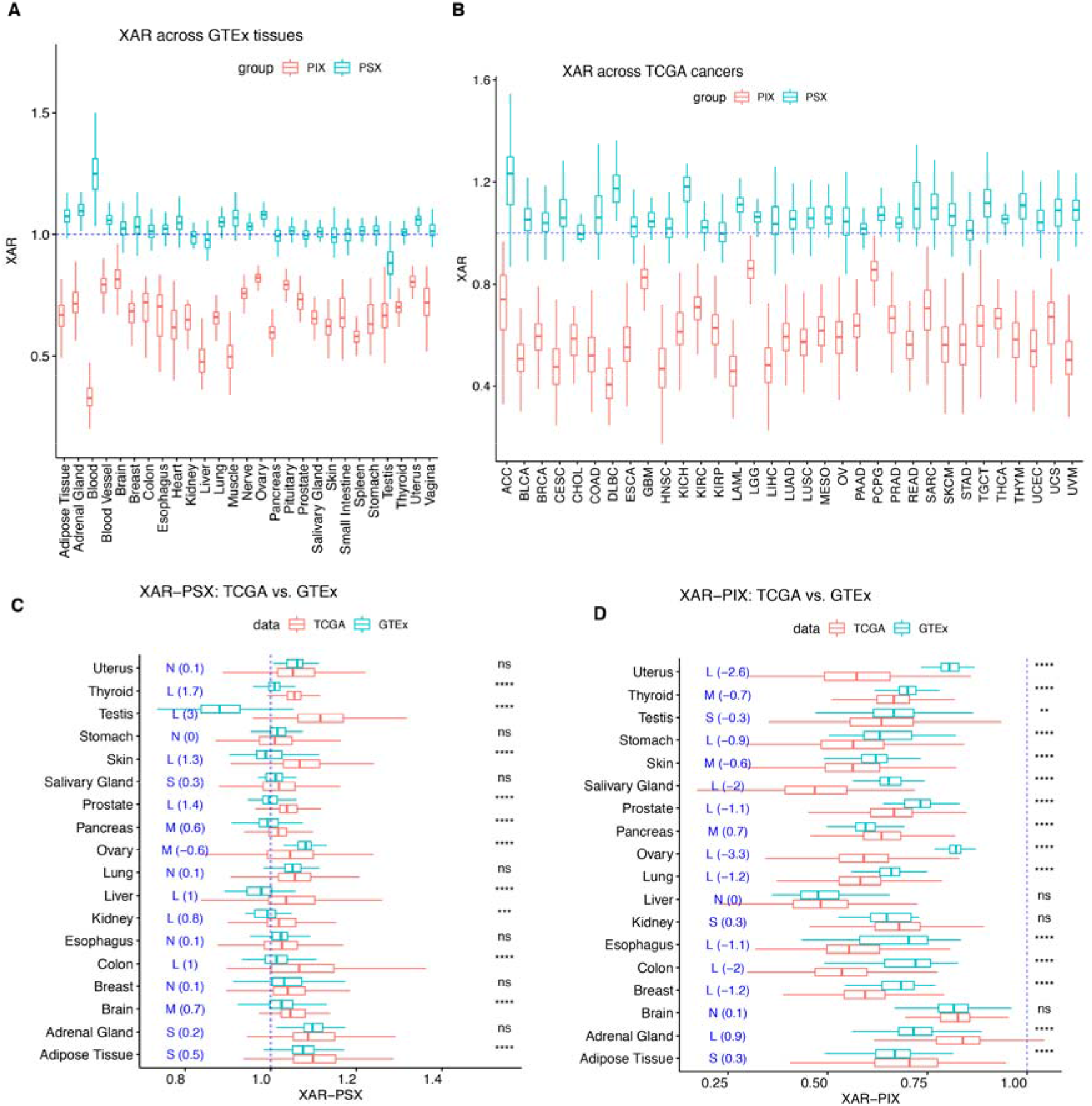
Differential dosages of PSX and PIX genes across normal tissues and cancer types. (**A**) XAR-PSX and XAR-PIX across 27 normal tissues from GTEx were close to one, except for blood and testis. (**B**) XAR-PSX and XAR-PIX across 33 cancer types from TCGA. (**C**) XAR-PSX were higher in cancers than in matched normal tissues. Effect sizes were indicated by Cohen’s D statistics in blue (negligible[N]<0.2, 0.2<=small[S]<0.5, 0.5<=moderate[M]<0.8, large[L]>=0.8). (**D**) XAR-PIX were lower in cancers than in matched normal tissues. Effect sizes were indicated by Cohen’s D statistics in blue. (**C-D**) Two-sided Wilcoxon test *P*-value significance: ns >0.05, * <= 0.05, ** <=0.01, *** <= 0.001, **** <= 0.0001. Matching of tumor types to normal tissues were as follows: ACC, Adrenal Gland; BRCA, Breast; COAD, Colon; ESCA, Esophagus; GBM, Brain; HNSC, Salivary Gland; KICH, Kidney; KIRC, Kidney; KIRP, Kidney; LGG, Brain; LIHC, Liver; LUAD, Lung; LUSC, Lung; OV, Ovary; PAAD, Pancreas; PRAD, Prostate; PCPG, Adrenal Gland; READ, Colon; SARC, Adipose Tissue; SKCM, Skin; STAD, Stomach; TGCT, Testis; THCA, Thyroid; UCS, Uterus; UCEC, Uterus. XAR: X-over-autosome dosage ratio, ACC: Adrenocortical carcinoma, BLCA: Bladder urothelial Carcinoma, BRCA: Breast invasive carcinoma, CESC: Cervical squamous cell carcinoma and endocervical adenocarcinoma, CHOL: Cholangiocarcinoma, COAD: Colon adenocarcinoma, CCRCC: Clear cell renal cell carcinoma, DLBC: Lymphoid neoplasm diffuse large B-cell lymphoma, ESCA: Esophageal carcinoma, GBM: Glioblastoma multiforme, HNSC: Head and neck squamous cell carcinoma, KICH: Kidney chromophobe, KIRC: Kidney renal clear cell carcinoma, KIRP: Kidney renal papillary cell carcinoma, LAML: Acute myeloid leukemia, LGG: Brain lower grade glioma, LIHC: Liver hepatocellular carcinoma, LUAD: Lung adenocarcinoma, LUSC: Lung squamous cell carcinoma, MESO: Mesothelioma, OV: Ovarian serous cystadenocarcinoma, PAAD: Pancreatic adenocarcinoma, PCPG: Pheochromocytoma and paraganglioma, PDA: Pancreatic ductal adenocarcinoma, PRAD: Prostate adenocarcinoma, READ: Rectum adenocarcinoma, SARC: Sarcoma, SKCM: Skin Cutaneous melanoma, STAD: Stomach adenocarcinoma, TGCT: Testicular germ cell tumors, THCA: Thyroid carcinoma, THYM: Thymoma, UCEC: Uterine corpus endometrial carcinoma, UCS: Uterine carcinosarcoma, UVM: Uveal melanoma.

Variations in TCGA cancers were generally larger than GTEx tissues for both XAR-PSX and XAR-PIX, reflecting abnormal dosage regulation across cancers. Surprisingly, across TCGA cancer types, we observed that XAR-PSX dosages were significantly increased (**Fig. 2C**), while XAR-PIX showed the opposite trend (**Fig. 2D**), compared to normal tissues from GTEx. Moreover, most magnitudes of changes (or effect sizes) of both XAR-PSX and XAR-PIX were moderate to large, as indicated by the Cohen’s D statistics. Of note, most tissues or cancer types showed no XAR difference between males and females, especially for XAR-PIX (**Fig. S3A-D**), which suggests that the possibility of XAR-PSX upregulation due to reactivation of the inactivated X in females (Chaligne, et al. 2015) can be excluded. Accordingly, when investigating proteomic data generated by mass spectrometry, the opposite trend between XAR-PIX and XAR-PSX was also observed when we compared primary tumors with normal tissue samples in CPTAC (**Fig. S2C-D**).

### XAR-PSX and XAR-PIX were differentially associated with stemness

It is well-known that XAR of stem cells changes dramatically during epigenetic reprogramming of early embryo development. On the other hand, increased stemness is an important feature or cancer cells (Malta, et al. 2018). We thus examined the relationship between XAR and stemness in cancer. We observed mostly positive correlations with cancer stemness for XAR-PSX, but mostly negative correlations for XAR-PIX, across cancer types (**Fig. 3A**). We further examined XAR in embryonic stem cells (ESC) and induced pluripotent stem cells (iPSC) using gene expression data from the Progenitor Cell Biology Consortium (PCBC). As expected, we found no difference between ESC and iPSC, for both XAR-PIX and XAR-PSX (**Fig. 3B**), probably because ESC and iPSC had similar expression pattern (Bock, et al. 2011). However, we found much lower XAR-PIX and XAR-PSX in male than in female iPSC (**Fig. 3C**), but the signals of sex difference in ESC were weak.

**Fig. 3.**
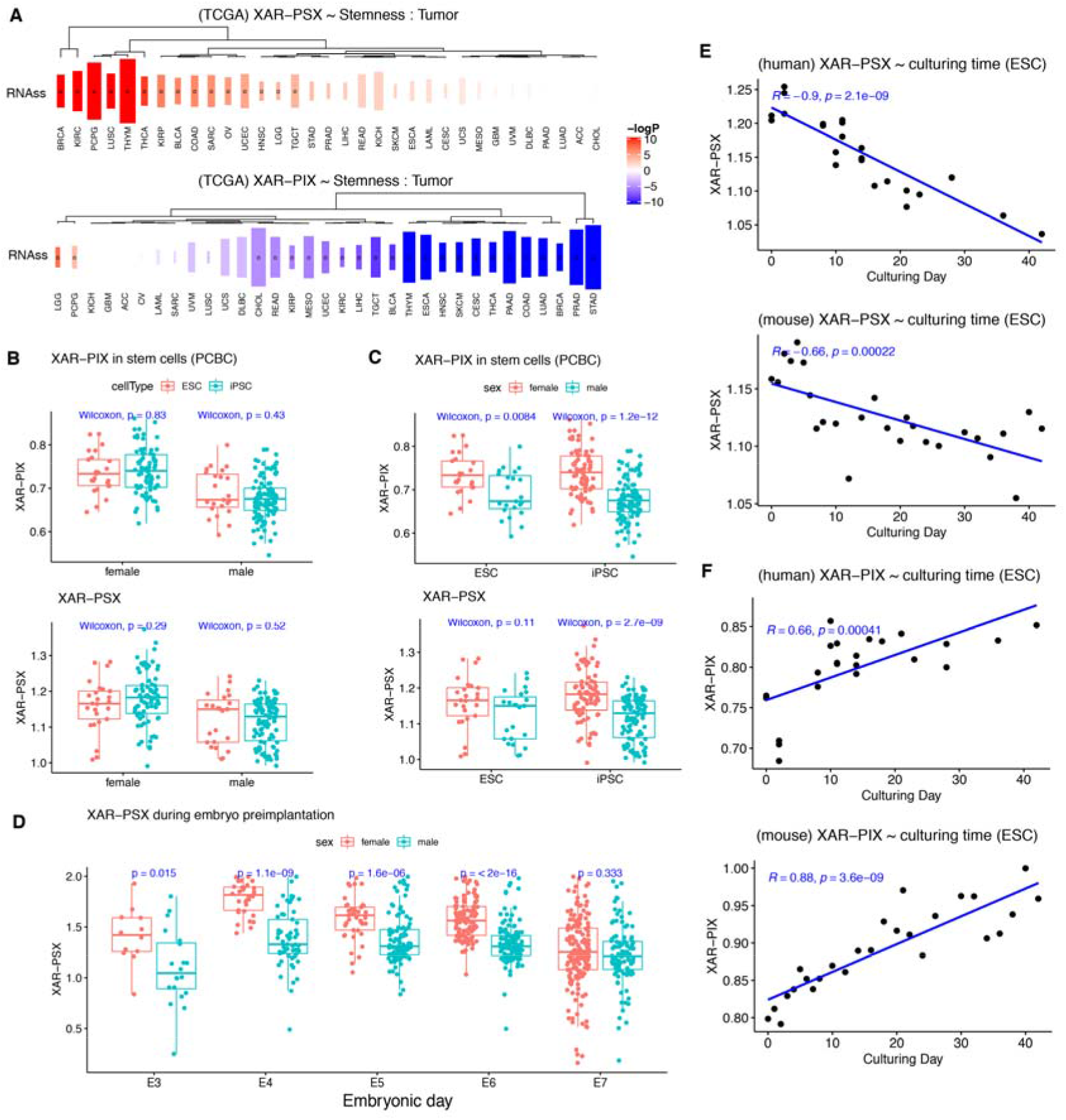
XAR-PSX and XAR-PIX were inversely associated with stemness. (**A**) Cancer stemness (RNAss) was positively correlated with XAR-PSX (top) and negatively with XAR-PIX (bottom). colors indicate -log10 P-values, and sizes of rectangles represent Pearson correlation. Correlations with FDR<0.1 were marked by circles. (**B**) Comparison of XAR-PIX (top) and XAR-PSX (bottom) between ESCs and iPSCs from females or males. (**C**) Comparison of XAR-PIX (top) and XAR-PSX (bottom) between sexes from ESCs or iPSCs. (**D**) XAR-PSX dynamics of embryonic stem cells (grouped by sex) during preimplantation from day 3 (E3) to 7 (E7). (E) Strong negative correlation of XAR-PSX with differentiation culturing time in both human (top) and mouse (bottom). (**F**) Strong positive correlation of XAR-PIX with differentiation culturing time in both human (top) and mouse (bottom). (**B-D**) Two-sided Wilcoxon test.

We noticed that XAR-PSX was between 1.1 to 1.2 in both ESCs and iPSCs, although ESCs and some iPSCs have two active X chromosomes (Bar, et al. 2019). This may be related to early embryonic development, during which XAR-PSX has peaked (as expected, close to 1.9 in female and 1.4 in male) at embryo day 4 (E4) and decreased dramatically to be equal in both sexes (around 1.2) (**Fig. 3D**), which was consistent with our observations of XAR-PSX in ESC and iPSC cell lines.

We further examined XAR dynamics during iPSC differentiation culturing. Interestingly, we found that XAR-PSX was reduced along iPSC culturing time (**Fig. 3E**), but XAR-PIX showed the opposite trend (**Fig. 3F**), in both human and mouse. These patterns were consistent with overall reduction of XAR-PIX and elevation of XAR-PSX across cancer types above.

### Insufficient PIX gene dosage in autoimmune patients

Both altered X-inactivation and X-linked risk genes were associated with SLE, we thus investigated which type of XAR (XAR-PIX or XAR-PSX) was dysregulated in SLE. By reanalyzing a previous dataset from SLE patients and healthy controls (Hung, et al. 2015), we found that XAR-PIX, but not XAR-PSX, were specifically down-regulated in SLE patients compared with healthy controls (**Fig. 4A**), which means insufficient dosage of X genes in SLE patients. Moreover, SLE patients with anti-Ro autoantibodies showed lower XAR-PIX dosage than those without anti-Ro autoantibodies (**Fig. 4B**). Similarly, SLE patients with more active interferon response (indicated by the Interferon Signature Metric, or ISM) showed lower XAR-PIX dosage (**Fig. 4C**). These findings may provide insights to understand SLE pathogenesis.

**Fig. 4.**
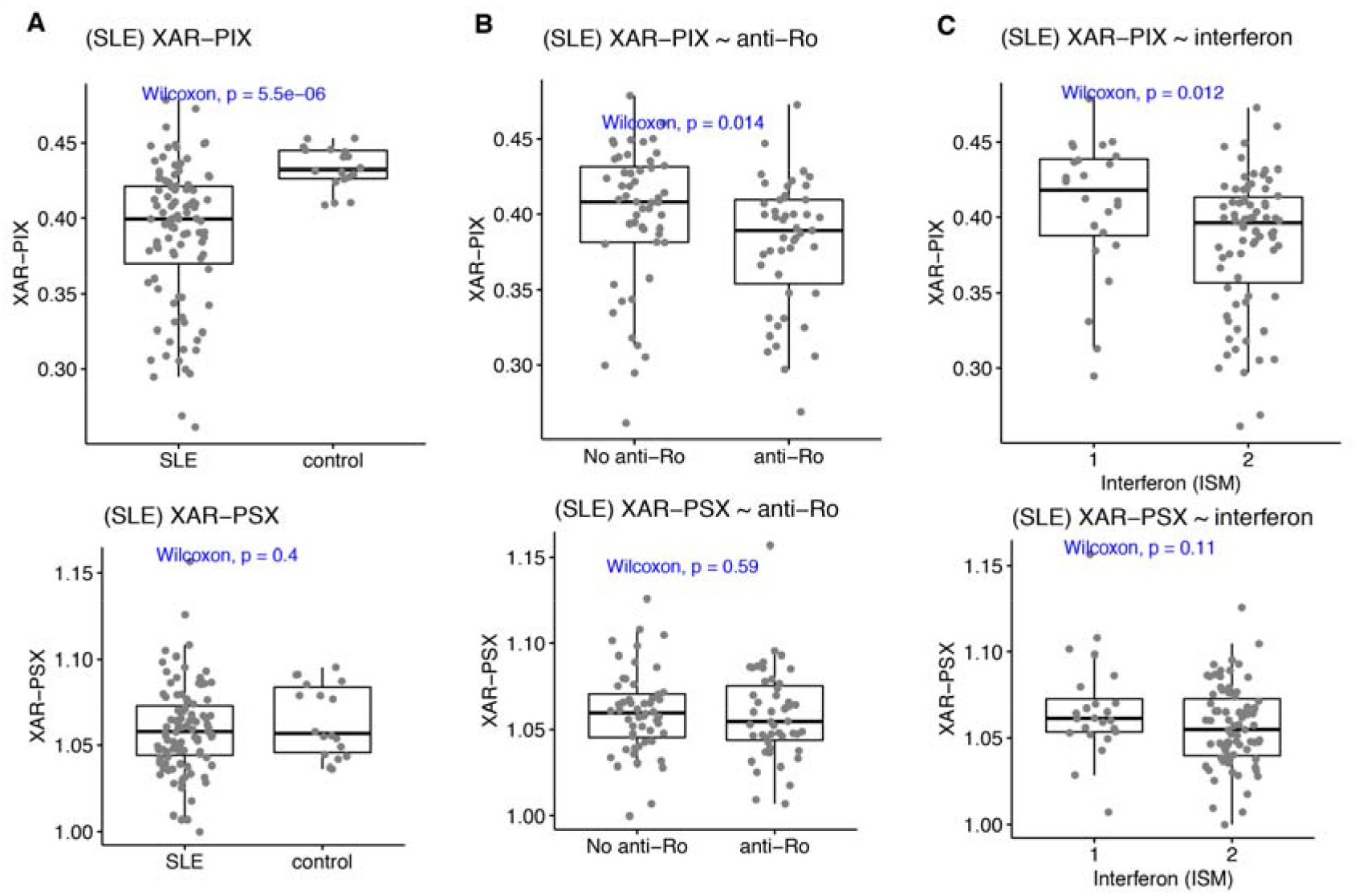
XAR in autoimmune disease. (**A**) XAR-PIX (top) and XAR-PSX (bottom) in SLE patients compared to healthy controls. (**B**) XAR-PIX (top) and XAR-PSX (bottom) in SLE patients with anti-Ro autoantibodies compared to those without anti-Ro. (**C**) XAR-PIX (top) and XAR-PSX (bottom) in SLE patients with more active interferon response (ISM) compared to those with less ISM. (**A-C**) Two-sided Wilcoxon test.

### TP53 and ATRX are new regulators of X dosage compensation

We found that only about 8.8% (29/329) and 10.5% (30/286) of PIX and PSX genes showed expression changes of more than 1.2-fold after knockdown of the key MSL component *KAT8* in mouse (Deng, et al. 2013). To identify new regulators of XAR, we examined TCGA mutation spectrums that may be associated with XAR in each cancer type. We reasoned that mutated XAR regulators should result in differential XAR compared with wildtype regulators. In total, we obtained 209 and 515 potential regulators (FDR < 0.5) for XAR-PIX and XAR-PSX in male tumors, respectively (**Fig. 5A, Fig. S4A**). For the top 200 regulators, gene ontology (GO) analysis was performed and similar GO terms were found for XAR-PIX and XAR-PSX regulators, mainly about post-translational modifications, including histone methylation, peptidyl-lysine methylation, and protein methylation (**Fig. S4B-C**).

**Fig. 5.**
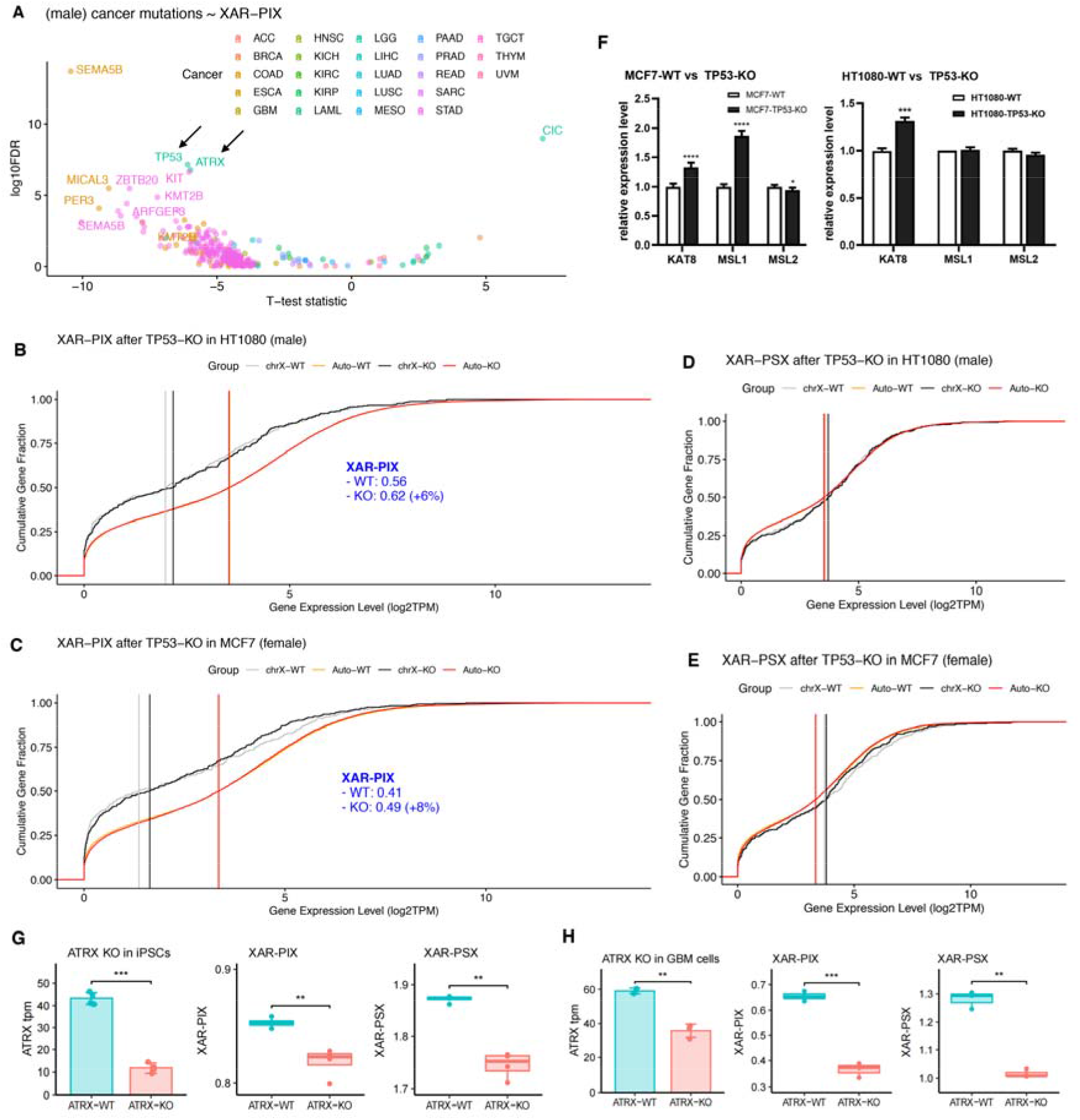
*TP53* and *ATRX* were two regulators of XAR. (**A**) Significance of XAR-PIX difference (y-axis, two-sided *t*-test) between wildtype and mutated genes across cancers in male. *TP53* and *ATRX* were marked by arrows. For better visualization, only genes with FDR<1 were shown as points and those with FDR<1e-4 labeled. (**B-C**) Cumulative distribution of PIX and autosomal gene expression levels for HT1080 (**B**) and MCF7 (**C**), before and after *TP53*-KO. Vertical lines indicate median expression used to calculate XAR-PIX dosages. (**D-E**) Cumulative distribution of PSX and autosomal gene expression levels for HT1080 (**D**) and MCF7 (**E**), before and after *TP53*-KO. Vertical lines indicate median expression used to calculate XAR-PSX dosages. (**F**) Gene expression changes of MSL complex subunits *KAT8, MSL1*, and *MSL2* in MCF7 (left) and HT1080 (right) cells, before and after *TP53*-KO. (**G**) *ATRX* knockout (KO) in mouse induced pluripotent stem cells (iPSCs) significantly reduced both XAR-PIX (middle) and XAR-PSX (right). (**H**) *ATRX* knockout (KO) in human GBM cells significantly reduced both XAR-PIX (middle) and XAR-PSX (right). (**F-H**) Two-sided *t*-test, * *P*<= 0.05, ** <=0.01, *** <= 0.001, **** <= 0.0001.

Among the potentially new regulators of XAR, we found that *TP53* and *ATRX* were promising regulators for XAR-PIX in both males and females (**Fig. 5A, Fig. S4D**), based on previous literatures mentioned above. They are both genome-wide chromatin remodelers, capable of regulating a large number of genes. We chose them for further investigation also for the following two reasons. We found that the top 200 XAR-PIX dependent genes formed a PPI network mostly centered on TP53 and ATRX (**Fig. S5**). TP53 was also a center node in the PPI network (**Fig. S4E**) with RNA-binding proteins (RBPs) associated with XAR-associated lncRNAs (Supplementary Results, **Fig. S4F, Table S2**).

Next, we constructed *TP53*-knockout (*TP53*-KO) cells from two *TP53*-wildtype (*TP53*-WT) cell lines, one male (HT1080) and one female (MCF7, containing three active X and no inactive X chromosomes (Sirchia, et al. 2005)), by using CRISPR technologies (**Fig. S4G, see Methods**), followed by RNA-seq. We observed XAR-PIX increase after *TP53*-KO in both sexes (**Fig. 5B and Fig. 5C**), in contrast to the unchanged XAR-PSX (**Fig. 5D and Fig. 5E**). We further demonstrated that *TP53* may partly regulate XAR-PIX by repressing KAT8, since *TP53*-KO enhanced *KAT8* expression in both sexes (**Fig. 5F**). *TP53*-KO had a larger effect in female cells (8% vs. 6% increase of XAR-PIX) than male cells, which may due to MCF7 cells containing three active X chromosomes and *TP53* affected *MSL1* in female cells only.

Finally, we examined the role of ATRX in XAR. We investigated XAR before and after *ATRX* knockout in two studies. Interestingly, ATRX-KO reduced both XAR-PIX and XAR-PSX in mouse iPSCs (**Fig. 5G**) and GBM cells (**Fig. 5H**). Similarly, we examined MSL expression changes after ATRX-KO, we observed consistent down-regulation of both MSL1 and MSL2 in either GBM or iPSC cells in mice, although KAT8 was found up-regulated in mouse iPSC cells while it was unchanged in GBM cells (**Fig. S6A**). To further support *ATRX* as a new regulator of XAR, we examined whether X-linked genes are more sensitive to the change of *ATRX* expression. For autosomal and X-linked genes co-expressed with *ATRX*, we compared their coefficient estimates of *ATRX* expression by linear regression models. We reasoned that larger absolute values of ATRX coefficients indicate higher impact on gene expression. As a result, *ATRX* regression coefficients were significantly higher in X-linked genes than in autosomal genes across a variety of tumors, at both mRNA (**Fig. S6B**) and protein level (**Fig. S6C**).

## Discussions

Here, we identified and comprehensively characterized two distinct groups of genes on the X chromosome. Integrating large-scale sequencing data from both cancer and normal samples greatly enhanced our ability to investigate both pathological and physiological roles of XAR. Our findings may help resolve XAR controversies described in our previous study (Xiong, et al. 2010). We summarized our findings in **Figure 6**. Our results showed that dosages of PIX and PSX genes were converging during stem cell differentiation but diverging in cancer cells. However, in SLE patients, only dosages of PIX genes, but not PSX genes, were down-regulated compared to healthy individuals. Finally, TP53 and ATRX were identified and validated as two XAR regulators.

**Fig. 6.**
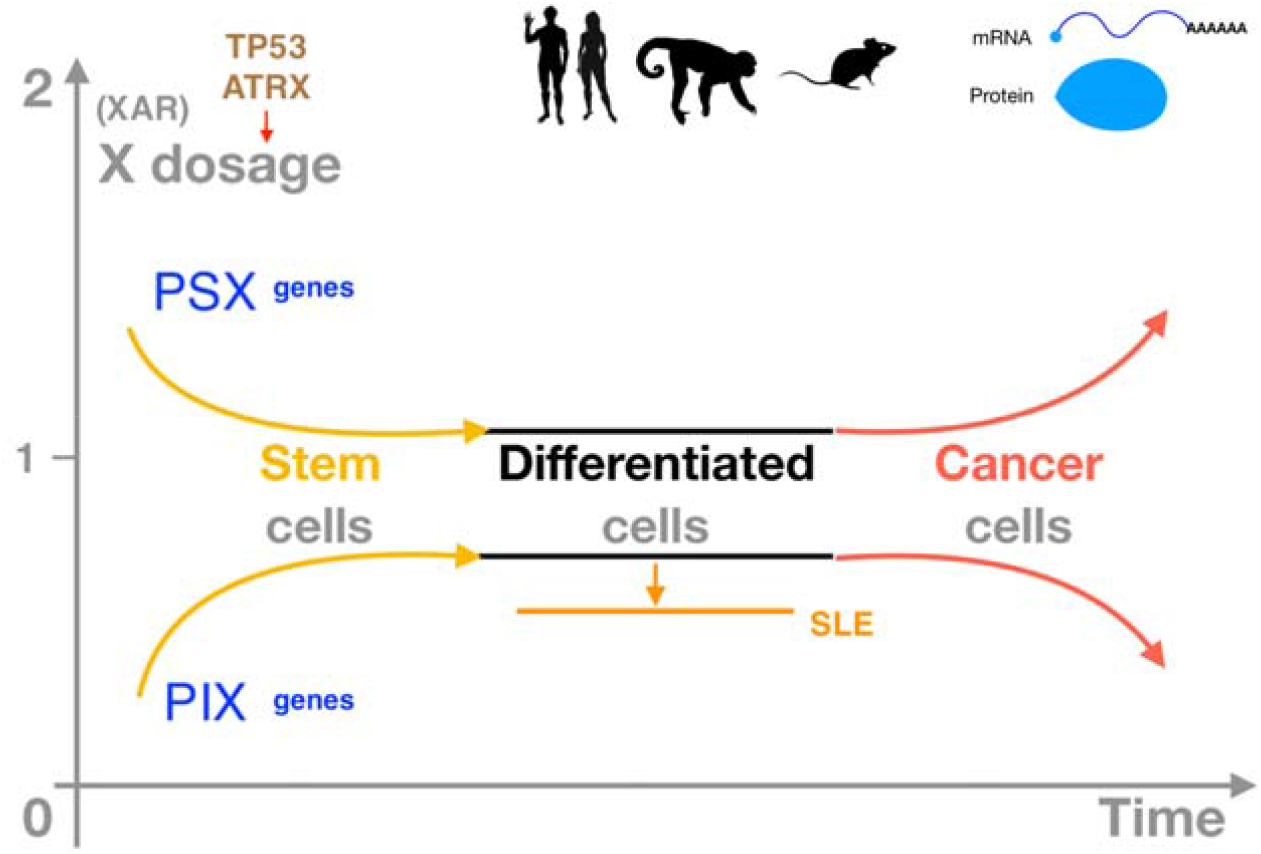
Major findings of this study. Differential dosages of two groups of X chromosome genes (PIX and PSX) were observed in human and animals at both mRNA and protein levels. X dosages were converging during stem cell differentiation but diverging in cancer. X dosage compensation was lower in blood cells of SLE autoimmune patients than in healthy individuals. TP53 and ATRX were identified and validated as two XAR regulators. XAR: X-over-autosome dosage ratio, PIX: ploidy-insensitive X genes, PSX: ploidy-sensitive X genes.

Although most genes from the TCEAL family, located within one of PIX gene clusters, were not intensively studied, most reports suggested that they acted as tumor suppressors, especially for *TCEAL7* (Lafferty-Whyte, et al. 2010; Yue, et al. 2019). For these tumor suppressors, lower sensitivity to dosage may protect both males and females from cancer equally. Interestingly, the chrX region around 100Mb showed the lowest expression level after zygote genome activation (ZGA) at E4 during embryo implantation (Petropoulos, et al. 2016).

Of note, *TP53* mRNA itself is a critical regulator of cellular stress (Haronikova, et al. 2019). A previous study found that the deadenylase poly(A)-specific ribonuclease (PARN) destabilized *TP53* mRNA while p53 increase activated PARN (Devany, et al. 2013), indicating a negative feedback loop between *TP53* mRNA and p53 protein levels. We observed that *TP53* mRNA was universally up-regulated in cancer samples, compared with non-cancerous samples, across cancer types (**Fig. S7**). Moreover, XAR-PIX was lower in tumors than non-cancerous samples across cancer types. These results were consistent with observations that p53 mutations (or higher *TP53* mRNA) were associated with lower XAR-PIX in cancer and *TP53*-KO resulted in XAR-PIX increase.

The frequency of transcriptional bursts and possible trans-acting enhancer-binding regulators were proposed to be the driving force behind X upregulation in mouse (Larsson, et al. 2019). Interestingly, p53 can bind pervasively to genome-wide enhancers (Younger and Rinn 2017) and regulate the transcriptional burst of target genes (Friedrich, et al. 2019), further supporting our finding of *TP53* regulating XAR. Moreover, the crosstalk between *TP53* and nonsense-mediated decay (NMD) pathway may contribute to XAR regulation by *TP53* (Gewandter, et al. 2011; Deng, et al. 2013). Therefore, *TP53* may regulate XAR by various mechanisms. We expect XAR regulation by *TP53* would be conserved among mammals, due to ultra-conservation of *TP53* during evolution (Lu, et al. 2009). However, further efforts are needed to support our hypothesis.

XAR in most cancers showed no difference between sexes, suggesting negligible contribution to XAR changes in cancer by reactivation of the inactivated X in females. Since *TP53* mutation-induced aneuploidies were not specific to X chromosome, they were unlikely to drive both XAR-PSX and XAR-PIX to change in such a coordinated way across cancers, as observed in this study. At least, XAR-PIX should be less affected by aneuploidies, by PIX definition.

Several limitations of this study represent directions for future research. First, it is unknown how distinct dosages of PIX and PSX genes were maintained and why did they show divergent trends in cancers compared to normal tissues. Second, despite the high cost and long period of CRISPR-based *TP53*-KO experiments, it would further solidify our conclusion if CRISPR-mediated *TP53*-KO can be established in more cell lines of different lineages. Third, additional XAR regulators remain to be identified by other ways, because this analysis was limited by mutation frequencies in cancer patients. Genome-wide screening experiments in the future probably will identify more XAR regulators. Until then, the problem of abnormal XAR-PIX and XAR-PSX in cancer may be solved.

## Materials and Methods

### Datasets

Gene expression levels (transcripts per million, or TPM) and related clinical information of RNA-seq samples from The Cancer Genome Atlas project (TCGA, https://portal.gdc.cancer.gov) and the Genotype-Tissue Expression project (GTEx, https://www.gtexportal.org), processed by the same computational pipeline, were obtained from the UCSC Xena database (https://xenabrowser.net) (Goldman, et al. 2020). For GTEx, tissues with less than 30 samples were excluded. Gene expression levels of cell lines sequenced by the Progenitor Cell Biology Consortium (PCBC) (Salomonis, et al. 2016) were downloaded from Synapse (https://www.synapse.org/#!Synapse:syn1773109/wiki/54962). Gene expression levels of ESCs during differentiation culturing for both human and mouse were downloaded from (Barry, et al. 2017). Somatic mutations for TCGA tumor samples were obtained the GDC Portal (https://portal.gdc.cancer.gov). Genome ploidy information for TCGA samples were downloaded from (Taylor, et al. 2018). Cancer stemness of TCGA samples were obtained from (Malta, et al. 2018). Processed single-cell RNA sequencing data for embryonic stem cells during preimplantation were downloaded from (Petropoulos, et al. 2016). Gene expression levels of SLE RNA-seq data were obtained from (Hung, et al. 2015). Half-life data of mRNAs were obtained from (Tani, et al. 2012). Our RNA-seq data for *TP53*-WT and *TP53*-KO cell lines (MCF7 and HT1080) from this study were deposited in the SRA database (PRJNA777384). Tissue-specific gene expression levels (Read counts) for non-human primates were downloaded from NHPRTR (http://nhprtr.org/) (Peng, et al. 2015). Gene expression levels (Fragments Per Kilobase of transcript per Million mapped reads, or FPKM) for the standard mouse strain, C57BL/6JJcl were collected from (Li, et al. 2017). Quantitative proteomic expression levels (TMT ratios) for tumors were obtained from CPTAC (https://pdc.cancer.gov/pdc/), covering data from 11 studies (PDC000120, PDC000116, PDC000110, PDC000127, PDC000204, PDC000221, PDC000153, PDC000234, PDC000270, PDC000125 and PDC000198). RNA-seq data for mouse ATRX knockout cell lines were collected from Gene Expression Omnibus (GEO, https://www.ncbi.nlm.nih.gov/geo/), including datasets GSE178113 (Qin, et al. 2022) and GSE107878 (Deneault, et al. 2018). LncRNA expression profiles for GTEx and TCGA samples were downloaded from RefLnc (http://reflnc.gao-lab.org/) (Jiang, et al. 2019).

### Defining PIX, PSX, and autosomal genes for dosage compensation estimation

For both X chromosome and autosomes, only protein-coding genes were considered and those with median expression levels TPM<=1 in GTEx samples were further excluded. For the remaining X chromosome genes, Pearson correlations (R) between gene expression and genome ploidy (Taylor, et al. 2018) were calculated in each cancer (except CHOL, DLBC, UCS, KICH, and BRCA) separately from TCGA, using only male samples to avoid the confounding of possible reactivation of the inactivated X in female samples. Then, the median of absolute R (absR) was calculated for each gene and the average of these absR medians (MR) was determined for these genes. Lastly, X chromosome genes with absR medians less or higher than MR were defined as PIX and PSX genes, respectively. Remaining PIX, PSX, and autosome genes were used for XAR-PIX and XAR-PSX calculation.

### Comparison of genomic features for PIX and PSX genes

Gene lengths were calculated for each gene either including or excluding introns, based on GENCODE v25 (Harrow, et al. 2012). PhastCons100way scores were downloaded from the UCSC Genome Browser (https://hgdownload.soe.ucsc.edu/downloads.html) and gene conservation were calculated by averaging phastCons100way scores within exonic regions.

GC content for each gene were calculated from its mRNA sequence as the percentage of G and C bases.

### Calculation of XAR-PSX and XAR-PIX

First, median expression levels of PSX, PIX, and autosomal genes were calculated. Then, XAR-PSX or XAR-PIX were defined as the ratio of PSX or PIX median over autosomal median, instead of the mean (Xiong, et al. 2010). The magnitude (Cohen’s D) of XAR difference between TCGA and GTEx was estimated by the function cohens_d from the R package rstatix (v0.6). For non-human species, genes were classified according to their homologous gene categories (PSX, PIX or autosomal genes) in humans, and then XAR-PSX and XAR-PIX were calculated as above, based on their processed gene expression levels (TPM). For proteomic expression levels, TMT ratios were used to calculate XAR, while XAR-PSX and XAR-PIX were compared between primary tumors and solid tissue normal samples.

### Association of XAR with somatic mutations

Damaging mutations were defined as Missense_Mutation, Nonsense_Mutation, Nonstop_Mutation, Splice_Region, Splice_Site, Translation_Start_Site, Frame_Shift_Del, Frame_Shift_Ins, In_Frame_Ins, or In_Frame_Del. For each gene, samples were grouped by whether having damaging mutations in each cancer for each gender. Genes with less than 20 damaging mutations were not considered. XAR between these two groups were compared using two-sided *t*-test and *P*-values were corrected using false discovery rate (FDR).

### Gene ontology analysis

GO enrichment analyses were performed using R package clusterProfiler (v4.2.2). Enriched biological processes were identified by right-tailed Fisher’s exact test, and Benjamini-Hochberg correction was used for multiple comparisons. For potential protein-coding regulators of XAR, genes expressed in at least in one tumor were used as the background gene set. For lncRNA analysis, neighboring protein-coding genes of XAR-associated lncRNAs were used as background gene set.

### ATRX regression coefficients comparison between autosomal and X-linked genes

For autosomal and X-linked genes, linear regression models with ATRX expression were fit at the mRNA or proteomic level in each tumor, in which *ATRX* coefficient estimates were collected and converted to absolute form for comparison between the two gene groups.

### Data visualization

The R package ggpubr (v.04, based on ggplot2) was used for basic data visualizations in this study. The heatmaps of stemness correlation with XAR were visualized by R package ComplexHeatmap (v2.2). Genomic distribution of PIX and PSX genes were visualized by R package circlize (v0.4). Correlation heatmaps between XAR predictions and observations were visualized by R package ggcorrplot2 (v0.1). Cumulative distribution of expression levels were visualized by R package ggplot2 (v3.3).

### Cell culture and CRISPR-Cas9 knockout

MCF7 and HT1080 cells were obtained from the American Type Culture Collection (ATCC). All cells were cultured in high-glucose DMEM containing 10% fetal bovine serum (FBS) and 1% penicillin/ streptomycin and maintained at 37 °C with 5% CO_2_.

The sgRNAs of *TP53* were designed on the CHOPCHOP website (https://chopchop.cbu.uib.no) as described before (Montague, et al. 2014). sgRNAs with highly predicted efficiency and lower off-target were chosen for in vitro transcription. The sgRNA sequence targeted *TP53* were as follows: sgRNA-1, 5’-TATCTGAGCAGCGCTCATGG-3’ and sgRNA-2, 5’-GGTGAGGCTCCCCTTTCTTG-3’. The sgRNA oligos were cloned into the pDR274 vector and in vitro transcribed using the MEGAshortscript T7 transcription kit (Thermo Fisher), then purifified using the MEGAclear kit (Thermo Fisher). The sgRNAs were then dissolved in RNase-free water and quantified using NanoDrop 1000 (Thermo Fisher).

To generate *TP53* knockout cells, electroporation of Cas9 RNP to the cells was performed as previously reported (Kim, et al. 2014). Specifically, 10 μg TrueCut Cas9 Protein v2 (Thermo) and 2 μg each of the in vitro transcribed sgRNAs were incubated together at 37°C for 20 min, and 5×10^5^ MCF7 or HT1080 cells (70%-90% confluent) were harvested and centrifuged at 90× g for 10 min. Then the Cas9 RNP was electroporated to the cell pellet by Lonza 4D-Nucleofector (Lonza) under a pre-optimized program.

### Western Blotting

Cells were harvested and lysed with RIPA buffer (Beyotime). Cell extracts of equal total protein (20 μg each) were separated by SDS-PAGE and transferred to Nylon membrane (Merck Millipore). The membranes were blocked with 5% nonfat milk in TBST washing buffer (10 mM Tris pH 8.0, 150 mM NaCl with 0.5% Tween-20) for 1 h at room temperature, then incubated at 4°C overnight with anti-*TP53* (1:1000; sc-126,Santa Cruz) and anti-GAPDH (1:5000; P30008M, Abmart) primary antibodies. After washing, the membranes were incubated with fluorescence-conjugated secondary antibodies and detected with Odyssey Imagers.

### RNA extraction and real-time quantitative PCR

Total RNA was extracted from cultured cells using RNAiso Plus (Takara) and reverse transcribed into cDNA by the PrimeScript™ RT reagent Kit (Takara), following the manufacturer’s protocol. Then real-time quantitative PCR was performed using TB Green® Premix Ex Taq™ II (Takara) on the Real-Time PCR Thermal Cycler qTOWER3 (Analytik Jena). Primers used in this study (forward/reverse; 5’ to 3’) were as follows:

*KAT8*: TCACTCGCAACCAAAAGCG/GATCGCCTCATGCTCCTTCT

*MSL1*: CCCATCACCGTTACCATTACG/GGAACAGCCAAGACTGAAGTTT

*MSL2*: GTAGCCACTGACTTATGTTCCAC/GCTGCAAATTAGGGCAACAGAC

GAPDH: GGAGCGAGATCCCTCCAAAAT/GGCTGTTGTCATACTTCTCATGG

Data analysis was performed using the 2(-ΔΔCT) method, and mRNA expression levels were normalized to GAPDH.

### RNA sequencing of MCF7 and HT1080 cell lines

Total RNA was extracted using TransZol Up Plus RNA Kit (Cat#ER501-01, Trans) following the manufacturer’s instructions and checked for RNA integrity by an Agilent Bioanalyzer 2100 (Agilent technologies, Santa Clara, CA, US). Qualified total RNA was further purified by RNAClean XP Kit (Cat#A63987, Beckman Coulter, Inc. Kraemer Boulevard Brea, CA,USA) and RNase-Free DNase Set (Cat#79254, QIAGEN, GmBH, Germany). Sequencing libraries were constructed following steps from the manufacturer’s instructions (VAHTS Universal V6 RNA-seq Library Prep Kit for Illumina®, Vazyme, NR604-02), followed by paired-end (2x 150bp) RNA-seq by Illumina HiSeq 2500. Cleaned reads were aligned to the human genome (hg38) by Kallisto (v0.46, default parameters) (Bray, et al. 2016) to obtain gene expression levels (TPM), based on gene annotations from GENCODE v25 (Harrow, et al. 2012). Reads were also mapped to hg38 using HISAT2 (v2.0.4, default parameters), and alignments of *TP53*-WT samples were visualized by IGV (v2.5.2) for manual inspection to confirm TP53 status.

## Data availability

The RNA-seq data generated in this study have been submitted to the NCBI BioProject database (https://www.ncbi.nlm.nih.gov/bioproject/) under accession number PRJNA777384.

## Acknowledgements

The work was supported by National Natural Science Foundation of China (NSFC) (Grant number 31571350, U1611265, and 31871323) and Guangdong Basic and Applied Basic Research Foundation (2021A1515110972). The authors would like to thank TCGA, GTEx, PCBC, NHPRTR, and other projects for making their research data publicly available.

## Author contributions

MBG and YYX conceived the project. MBG and ZWF collected the data, performed the analysis, and interpreted the results. MBG wrote the manuscript with input from ZWF. ZWF and BHC performed the experimental validation. ZSY was involved in project discussion. YYX was involved in critical discussions and reviewed the manuscript. All authors approved the manuscript.

## References

Allison SJ, Milner J. 2004. Remodelling chromatin on a global scale: a novel protective function of p53. Carcinogenesis 25:1551–1557.

Bar S, Seaton LR, Weissbein U, Eldar-Geva T, Benvenisty N. 2019. Global Characterization of X Chromosome Inactivation in Human Pluripotent Stem Cells. Cell Rep 27:20–29 e23.

Barry C, Schmitz MT, Jiang P, Schwartz MP, Duffin BM, Swanson S, Bacher R, Bolin JM, Elwell AL, McIntosh BE, et al. 2017. Species-specific developmental timing is maintained by pluripotent stem cells ex utero. Dev Biol 423:101–110.

Bock C, Kiskinis E, Verstappen G, Gu H, Boulting G, Smith ZD, Ziller M, Croft GF, Amoroso MW, Oakley DH, et al. 2011. Reference Maps of human ES and iPS cell variation enable high-throughput characterization of pluripotent cell lines. Cell 144:439–452.

Bray NL, Pimentel H, Melsted P, Pachter L. 2016. Near-optimal probabilistic RNA-seq quantification. Nat Biotechnol 34:525–527.

Chaligne R, Popova T, Mendoza-Parra MA, Saleem MA, Gentien D, Ban K, Piolot T, Leroy O, Mariani O, Gronemeyer H, et al. 2015. The inactive X chromosome is epigenetically unstable and transcriptionally labile in breast cancer. Genome Res 25:488–503.

Courel M, Clement Y, Bossevain C, Foretek D, Vidal Cruchez O, Yi Z, Benard M, Benassy MN, Kress M, Vindry C, et al. 2019. GC content shapes mRNA storage and decay in human cells. Elife 8.

Davoli T, Xu AW, Mengwasser KE, Sack LM, Yoon JC, Park PJ, Elledge SJ. 2013. Cumulative haploinsufficiency and triplosensitivity drive aneuploidy patterns and shape the cancer genome. Cell 155:948–962.

Deneault E, White SH, Rodrigues DC, Ross PJ, Faheem M, Zaslavsky K, Wang Z, Alexandrova R, Pellecchia G, Wei W, et al. 2018. Complete Disruption of Autism-Susceptibility Genes by Gene Editing Predominantly Reduces Functional Connectivity of Isogenic Human Neurons. Stem Cell Reports 11:1211–1225.

Deng X, Berletch JB, Ma W, Nguyen DK, Hiatt JB, Noble WS, Shendure J, Disteche CM. 2013. Mammalian X upregulation is associated with enhanced transcription initiation, RNA half-life, and MOF-mediated H4K16 acetylation. Dev Cell 25:55–68.

Devany E, Zhang X, Park JY, Tian B, Kleiman FE. 2013. Positive and negative feedback loops in the p53 and mRNA 3’ processing pathways. Proc Natl Acad Sci U S A 110:3351–3356.

Disteche CM. 2012. Dosage compensation of the sex chromosomes. Annu Rev Genet 46:537–560.

Donehower LA, Soussi T, Korkut A, Liu Y, Schultz A, Cardenas M, Li X, Babur O, Hsu TK, Lichtarge O, et al. 2019. Integrated Analysis of TP53 Gene and Pathway Alterations in The Cancer Genome Atlas. Cell Rep 28:3010.

Dunford A, Weinstock DM, Savova V, Schumacher SE, Cleary JP, Yoda A, Sullivan TJ, Hess JM, Gimelbrant AA, Beroukhim R, et al. 2017. Tumor-suppressor genes that escape from X-inactivation contribute to cancer sex bias. Nat Genet 49:10–16.

Dyer MA, Qadeer ZA, Valle-Garcia D, Bernstein E. 2017. ATRX and DAXX: Mechanisms and Mutations. Cold Spring Harb Perspect Med 7.

Friedrich D, Friedel L, Finzel A, Herrmann A, Preibisch S, Loewer A. 2019. Stochastic transcription in the p53-mediated response to DNA damage is modulated by burst frequency. Mol Syst Biol 15:e9068.

Garrick D, Sharpe JA, Arkell R, Dobbie L, Smith AJ, Wood WG, Higgs DR, Gibbons RJ. 2006. Loss of Atrx affects trophoblast development and the pattern of X-inactivation in extraembryonic tissues. PLoS Genet 2:e58.

Gewandter JS, Bambara RA, O’Reilly MA. 2011. The RNA surveillance protein SMG1 activates p53 in response to DNA double-strand breaks but not exogenously oxidized mRNA. Cell Cycle 10:2561–2567.

Goldman MJ, Craft B, Hastie M, Repecka K, McDade F, Kamath A, Banerjee A, Luo Y, Rogers D, Brooks AN, et al. 2020. Visualizing and interpreting cancer genomics data via the Xena platform. Nat Biotechnol 38:675–678.

GTEx-Consortium. 2013. The Genotype-Tissue Expression (GTEx) project. Nat Genet 45:580–585.

Gulve N, Su C, Deng Z, Soldan SS, Vladimirova O, Wickramasinghe J, Zheng H, Kossenkov AV, Lieberman PM. 2022. DAXX-ATRX regulation of p53 chromatin binding and DNA damage response. Nat Commun 13:5033.

Hainaut P, Pfeifer GP. 2016. Somatic TP53 Mutations in the Era of Genome Sequencing. Cold Spring Harb Perspect Med 6.

Hall LL, Byron M, Butler J, Becker KA, Nelson A, Amit M, Itskovitz-Eldor J, Stein J, Stein G, Ware C, et al. 2008. X-inactivation reveals epigenetic anomalies in most hESC but identifies sublines that initiate as expected. J Cell Physiol 216:445–452.

Haronikova L, Olivares-Illana V, Wang L, Karakostis K, Chen S, Fahraeus R. 2019. The p53 mRNA: an integral part of the cellular stress response. Nucleic Acids Res 47:3257–3271.

Harrow J, Frankish A, Gonzalez JM, Tapanari E, Diekhans M, Kokocinski F, Aken BL, Barrell D, Zadissa A, Searle S, et al. 2012. GENCODE: the reference human genome annotation for The ENCODE Project. Genome Res 22:1760–1774.

Haupt S, Caramia F, Herschtal A, Soussi T, Lozano G, Chen H, Liang H, Speed TP, Haupt Y. 2019. Identification of cancer sex-disparity in the functional integrity of p53 and its X chromosome network. Nat Commun 10:5385.

He Q, Kim H, Huang R, Lu W, Tang M, Shi F, Yang D, Zhang X, Huang J, Liu D, et al. 2015. The Daxx/Atrx Complex Protects Tandem Repetitive Elements during DNA Hypomethylation by Promoting H3K9 Trimethylation. Cell Stem Cell 17:273–286.

Hewagama A, Gorelik G, Patel D, Liyanarachchi P, McCune WJ, Somers E, Gonzalez-Rivera T, Michigan Lupus C, Strickland F, Richardson B. 2013. Overexpression of X-linked genes in T cells from women with lupus. J Autoimmun 41:60–71.

Hu Y, Shi G, Zhang L, Li F, Jiang Y, Jiang S, Ma W, Zhao Y, Songyang Z, Huang J. 2016. Switch telomerase to ALT mechanism by inducing telomeric DNA damages and dysfunction of ATRX and DAXX. Sci Rep 6:32280.

Hung T, Pratt GA, Sundararaman B, Townsend MJ, Chaivorapol C, Bhangale T, Graham RR, Ortmann W, Criswell LA, Yeo GW, et al. 2015. The Ro60 autoantigen binds endogenous retroelements and regulates inflammatory gene expression. Science 350:455–459.

Jiang S, Cheng SJ, Ren LC, Wang Q, Kang YJ, Ding Y, Hou M, Yang XX, Lin Y, Liang N, et al. 2019. An expanded landscape of human long noncoding RNA. Nucleic Acids Res 47:7842–7856.

Julien P, Brawand D, Soumillon M, Necsulea A, Liechti A, Schutz F, Daish T, Grutzner F, Kaessmann H. 2012. Mechanisms and evolutionary patterns of mammalian and avian dosage compensation. PLoS Biol 10:e1001328.

Kim S, Kim D, Cho SW, Kim J, Kim JS. 2014. Highly efficient RNA-guided genome editing in human cells via delivery of purified Cas9 ribonucleoproteins. Genome Res 24:1012–1019.

Kruse JP, Gu W. 2009. MSL2 promotes Mdm2-independent cytoplasmic localization of p53. J Biol Chem 284:3250–3263.

Lafferty-Whyte K, Bilsland A, Hoare SF, Burns S, Zaffaroni N, Cairney CJ, Keith WN. 2010. TCEAL7 inhibition of c-Myc activity in alternative lengthening of telomeres regulates hTERT expression. Neoplasia 12:405–414.

Larsson AJM, Coucoravas C, Sandberg R, Reinius B. 2019. X-chromosome upregulation is driven by increased burst frequency. Nat Struct Mol Biol 26:963–969.

Li B, Qing T, Zhu J, Wen Z, Yu Y, Fukumura R, Zheng Y, Gondo Y, Shi L. 2017. A Comprehensive Mouse Transcriptomic BodyMap across 17 Tissues by RNA-seq. Sci Rep 7:4200.

Li X, Wu L, Corsa CA, Kunkel S, Dou Y. 2009. Two mammalian MOF complexes regulate transcription activation by distinct mechanisms. Mol Cell 36:290–301.

Linke SP, Sengupta S, Khabie N, Jeffries BA, Buchhop S, Miska S, Henning W, Pedeux R, Wang XW, Hofseth LJ, et al. 2003. p53 interacts with hRAD51 and hRAD54, and directly modulates homologous recombination. Cancer Res 63:2596–2605.

Lu WJ, Amatruda JF, Abrams JM. 2009. p53 ancestry: gazing through an evolutionary lens. Nat Rev Cancer 9:758–762.

Malta TM, Sokolov A, Gentles AJ, Burzykowski T, Poisson L, Weinstein JN, Kaminska B, Huelsken J, Omberg L, Gevaert O, et al. 2018. Machine Learning Identifies Stemness Features Associated with Oncogenic Dedifferentiation. Cell 173:338–354 e315.

Meisel RP, Connallon T. 2013. The faster-X effect: integrating theory and data. Trends Genet 29:537–544.

Montague TG, Cruz JM, Gagnon JA, Church GM, Valen E. 2014. CHOPCHOP: a CRISPR/Cas9 and TALEN web tool for genome editing. Nucleic Acids Res 42:W401–407.

Munoz-Fontela C, Mandinova A, Aaronson SA, Lee SW. 2016. Emerging roles of p53 and other tumour-suppressor genes in immune regulation. Nat Rev Immunol 16:741–750.

Nguyen TT, Grimm SA, Bushel PR, Li J, Li Y, Bennett BD, Lavender CA, Ward JM, Fargo DC, Anderson CW, et al. 2018. Revealing a human p53 universe. Nucleic Acids Res 46:8153–8167.

Oppel F, Tao T, Shi H, Ross KN, Zimmerman MW, He S, Tong G, Aster JC, Look AT. 2019. Loss of atrx cooperates with p53-deficiency to promote the development of sarcomas and other malignancies. PLoS Genet 15:e1008039.

Pageau GJ, Hall LL, Ganesan S, Livingston DM, Lawrence JB. 2007. The disappearing Barr body in breast and ovarian cancers. Nat Rev Cancer 7:628–633.

Peng X, Thierry-Mieg J, Thierry-Mieg D, Nishida A, Pipes L, Bozinoski M, Thomas MJ, Kelly S, Weiss JM, Raveendran M, et al. 2015. Tissue-specific transcriptome sequencing analysis expands the non-human primate reference transcriptome resource (NHPRTR). Nucleic Acids Res 43:D737–742.

Petropoulos S, Edsgard D, Reinius B, Deng Q, Panula SP, Codeluppi S, Plaza Reyes A, Linnarsson S, Sandberg R, Lanner F. 2016. Single-Cell RNA-Seq Reveals Lineage and X Chromosome Dynamics in Human Preimplantation Embryos. Cell 165:1012–1026.

Qin T, Mullan B, Ravindran R, Messinger D, Siada R, Cummings JR, Harris M, Muruganand A, Pyaram K, Miklja Z, et al. 2022. ATRX loss in glioma results in dysregulation of cell-cycle phase transition and ATM inhibitor radio-sensitization. Cell Rep 38:110216.

Ren W, Medeiros N, Warneford-Thomson R, Wulfridge P, Yan Q, Bian J, Sidoli S, Garcia BA, Skordalakes E, Joyce E, et al. 2020. Disruption of ATRX-RNA interactions uncovers roles in ATRX localization and PRC2 function. Nat Commun 11:2219.

Salomonis N, Dexheimer PJ, Omberg L, Schroll R, Bush S, Huo J, Schriml L, Ho Sui S, Keddache M, Mayhew C, et al. 2016. Integrated Genomic Analysis of Diverse Induced Pluripotent Stem Cells from the Progenitor Cell Biology Consortium. Stem Cell Reports 7:110–125.

Seah C, Levy MA, Jiang Y, Mokhtarzada S, Higgs DR, Gibbons RJ, Berube NG. 2008. Neuronal death resulting from targeted disruption of the Snf2 protein ATRX is mediated by p53. J Neurosci 28:12570–12580.

Sirchia SM, Ramoscelli L, Grati FR, Barbera F, Coradini D, Rossella F, Porta G, Lesma E, Ruggeri A, Radice P, et al. 2005. Loss of the inactive X chromosome and replication of the active X in BRCA1-defective and wild-type breast cancer cells. Cancer Res 65:2139–2146.

Syrett CM, Paneru B, Sandoval-Heglund D, Wang J, Banerjee S, Sindhava V, Behrens EM, Atchison M, Anguera MC. 2019. Altered X-chromosome inactivation in T cells may promote sex-biased autoimmune diseases. JCI Insight 4.

Tani H, Mizutani R, Salam KA, Tano K, Ijiri K, Wakamatsu A, Isogai T, Suzuki Y, Akimitsu N. 2012. Genome-wide determination of RNA stability reveals hundreds of short-lived noncoding transcripts in mammals. Genome Res 22:947–956.

Taylor AM, Shih J, Ha G, Gao GF, Zhang X, Berger AC, Schumacher SE, Wang C, Hu H, Liu J, et al. 2018. Genomic and Functional Approaches to Understanding Cancer Aneuploidy. Cancer Cell 33:676–689 e673.

Teng YC, Sundaresan A, O’Hara R, Gant VU, Li M, Martire S, Warshaw JN, Basu A, Banaszynski LA. 2021. ATRX promotes heterochromatin formation to protect cells from G-quadruplex DNA-mediated stress. Nat Commun 12:3887.

Vallot C, Ouimette JF, Rougeulle C. 2016. Establishment of X chromosome inactivation and epigenomic features of the inactive X depend on cellular contexts. Bioessays 38:869–880.

Xiong Y, Chen X, Chen Z, Wang X, Shi S, Wang X, Zhang J, He X. 2010. RNA sequencing shows no dosage compensation of the active X-chromosome. Nat Genet 42:1043–1047.

Yang L, Yildirim E, Kirby JE, Press W, Lee JT. 2020. Widespread organ tolerance to Xist loss and X reactivation except under chronic stress in the gut. Proc Natl Acad Sci U S A 117:4262–4272.

Younger ST, Rinn JL. 2017. p53 regulates enhancer accessibility and activity in response to DNA damage. Nucleic Acids Res 45:9889–9900.

Yue X, Lan F, Xia T. 2019. Hypoxic Glioma Cell-Secreted Exosomal miR-301a Activates Wnt/beta-catenin Signaling and Promotes Radiation Resistance by Targeting TCEAL7. Mol Ther 27:1939–1949.

