## Supplementary Figures S1-S7 for "Comprehensively characterizing evolutionarily conserved differential dosage compensations of the X chromosome during development and in diseases"

**Supplementary Results**

**Potential lncRNA regulators of XAR**

LncRNAs regulate gene expression at multiple levels, including chromatin remodeling, transcription and post-transcription regulation (Statello, et al. 2021). One of the best-known examples of gene expression regulation by lncRNA is X inactive-specific transcript (*XIST*)-mediated X inactivation in females (Loda, et al. 2022). Considering the important roles of lncRNAs in gene expression regulation, we speculated that there were other lncRNAs related to dosage compensation to be explored. Therefore, we correlated lncRNA with XAR in each cancer in male, and obtained 2,689 and 31 lncRNAs significantly related to XAR-PIX and XAR-PSX, respectively. Next, we focused on the co-expressed genes and RNA-protein interactions of these lncRNAs, which might provide clues for how these lncRNAs affect dosage compensation in tumors. As a results, co-expressed genes of these lncRNAs were most enriched in histone modification (**Fig. S4F**), consistent with the results in potential protein-coding regulators of XAR (**Fig. S4B-C**), suggesting that lncRNAs might involve in dosage compensation by regulating genes related to histone modification. In addition, genes functioning in nuclear transport, chromatin organization, and mRNA metabolic processes were enriched. In terms of predicted interactions with XAR-associated lncRNAs, proteins involved in RNA processing and stability as well as translational regulation were enriched (**Table S2**).

**Supplementary Methods**

**Correlations for lncRNAs and XAR in tumors**

For 77,900 lncRNAs collected, those with median TPM greater than 0 in at least 1 tumor were selected. Next, linear regression models were fit to lncRNAs expression and XAR in each tumor, with lncRNAs correlated to XAR-PIX or XAR-PSX in at least 3 male tumors extracted for further study. Neighboring co-expressing protein-coding genes of these lncRNAs were identified by Pearson correlation coefficient (FDR<0.05) and subjected to GO analysis. RNA-protein interactions of XAR-associated lncRNAs were collected from LncSEA2.0 (<http://bio.liclab.net/LncSEA/index.php>).

**Supplementary Figures**

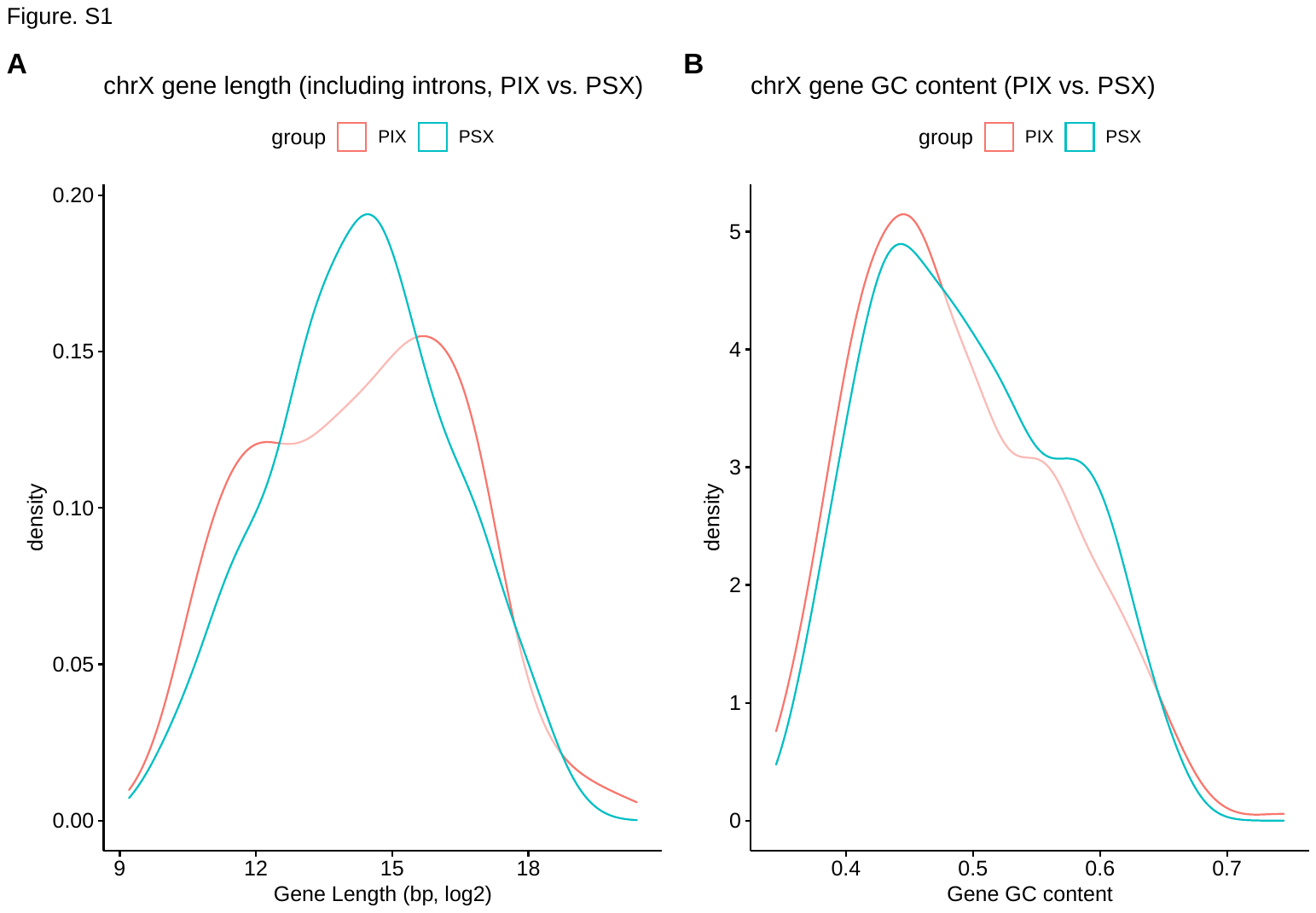

**Fig. S1,** (**A-B**) Comparing gene length (including introns, **A**) and GC content (**B**) between PIX and PSX genes.

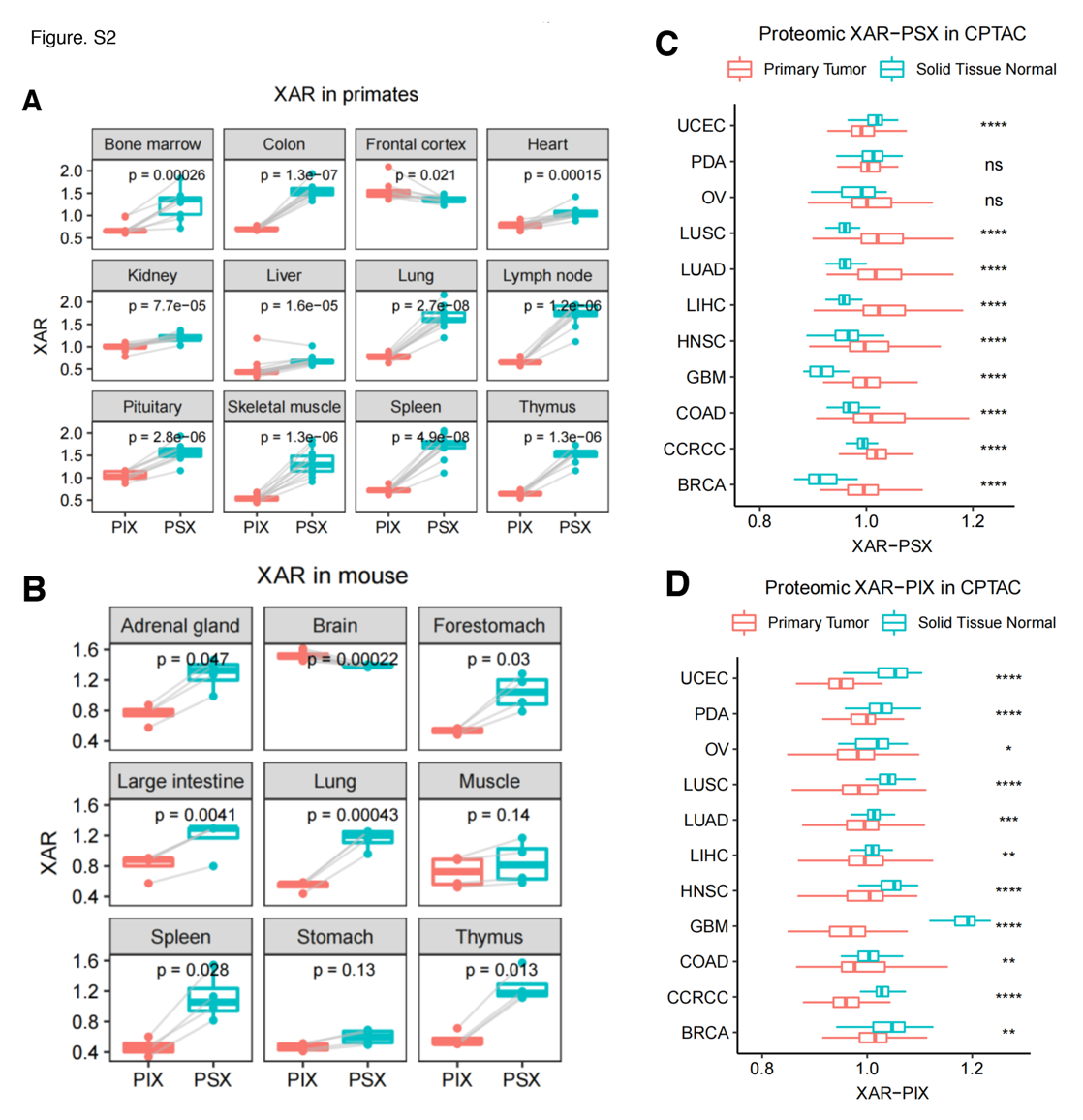

**Fig. S2, Differential XAR-PSX and XAR-PIX at mRNA level in animals and at protein level in human.**

(**A**) Evolutionarily conserved differential dosages between PIX and PSX genes in primates across 12 tissues. (**B**) Evolutionarily conserved differential dosages between PIX and PSX genes in mouse across 9 tissues. (**C**) Protein-level XAR-PSX were higher in tumors than normal tissues in most cancer types. (**D**) Protein-level XAR-PIX were lower in tumors than normal tissues of all investigated cancer types.

(**A-B**) Two-sided paired t-test. (**C-D**) Two-sided Wilcoxon test *P*-value significance: ns >0.05, * <= 0.05, ** <=0.01, *** <= 0.001, **** <= 0.0001.

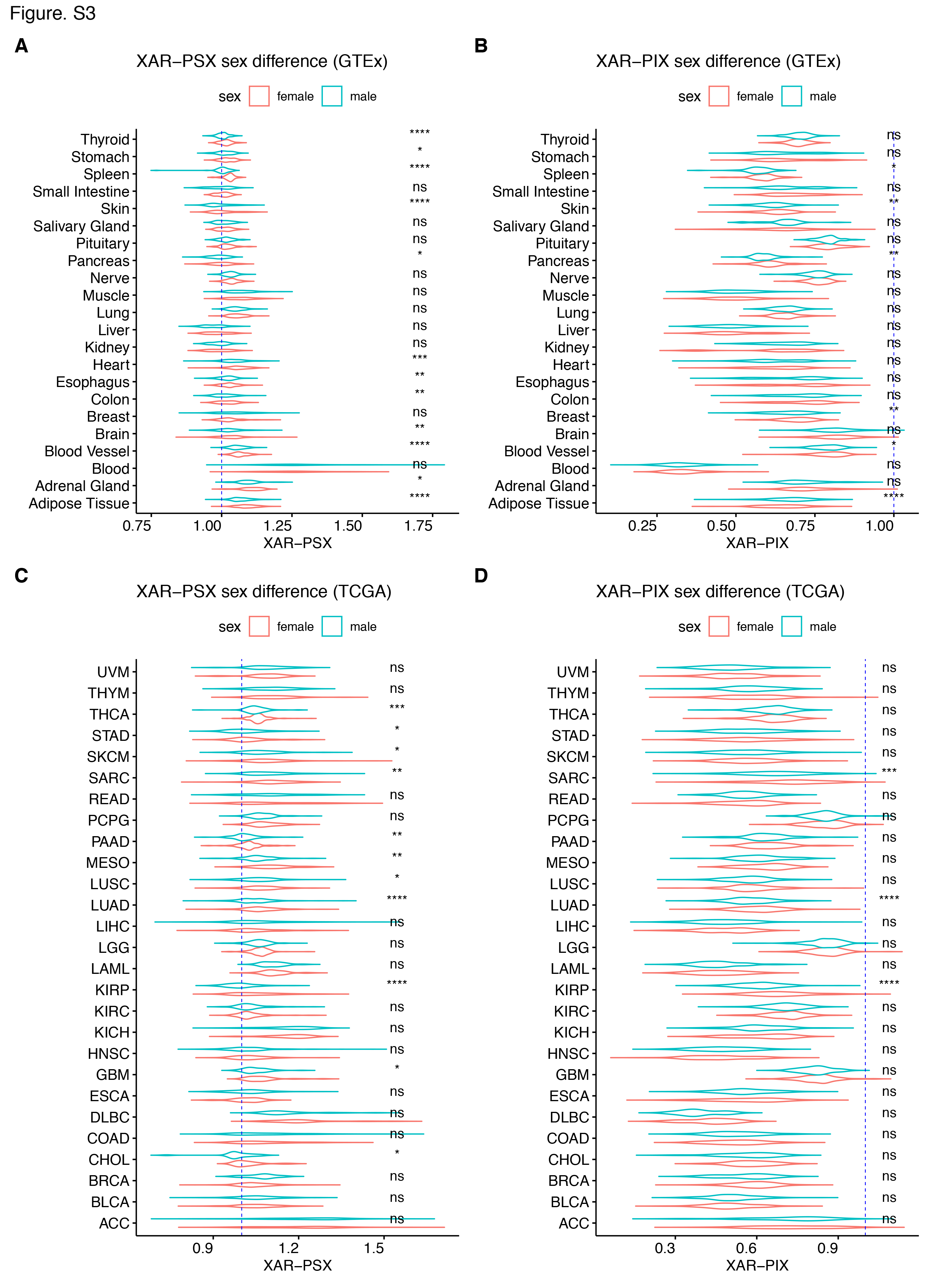

**Fig. S3,** **Sex difference of XAR**. (**A-B**) Comparison of XAR-PSX (**A**) and XAR-PIX (**B**) in GTEx normal tissues. (**C-D**) Comparison of XAR-PSX (**C**) and XAR-PIX (**D**) in TCGA cancer types. Wilcoxon test, *P*-value significance: ns >0.05, * <= 0.05, ** <=0.01, *** <= 0.001, **** <= 0.0001.

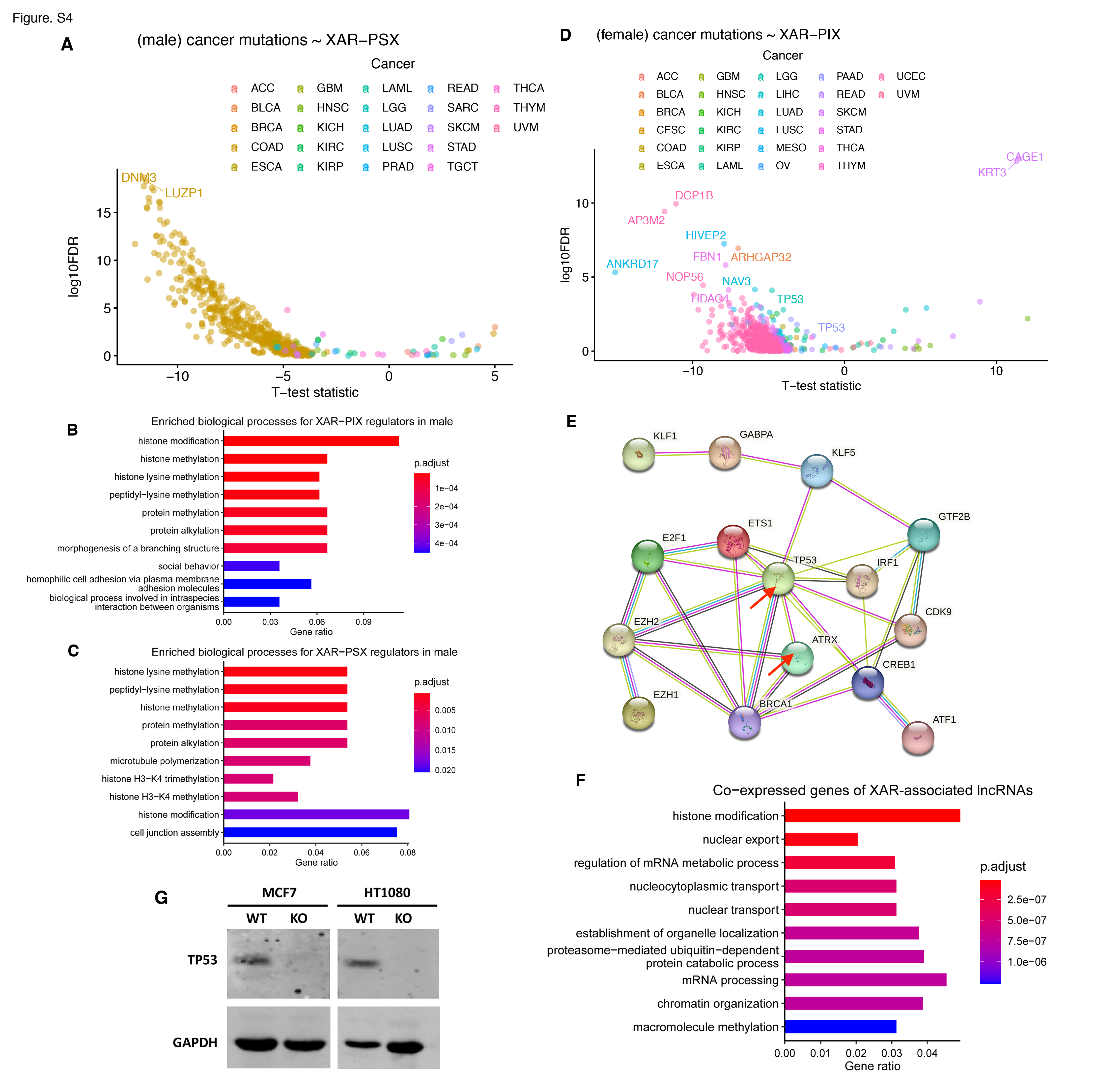

**Fig. S4, Potential XAR regulators revealed by TCGA mutation analysis**. (**A**) Significance of XAR-PSX difference (y-axis, two-sided *t*-test) between wildtype and mutated genes across cancers in male. For better visualization, only genes with FDR<1 were shown as points and those with FDR<1e-4 labeled. (**B**-**C**) Top 10 enriched biological processes in GO analysis for top 200 potential regulators of XAR-PIX (**B**) and XAR-PSX (**C**). (**D**) Western blot results for CRISPR-based *TP53*-KO in MCF7 (female) and HT1080 (male) cells. (**D**) Significance of XAR-PIX difference (y-axis, two-sided *t*-test) between wildtype and mutated genes across cancers in female. (**E**) The PPI network of TP53, ATRX, and RBPs enriched in XAR-associated lncRNAs. (**F**) Top 10 enriched biological processes for co-expressed genes of XAR-associated lnRNAs. (**G**) Western blot results for CRISPR-based *TP53*-KO in MCF7 (female) and HT1080 (male) cells.

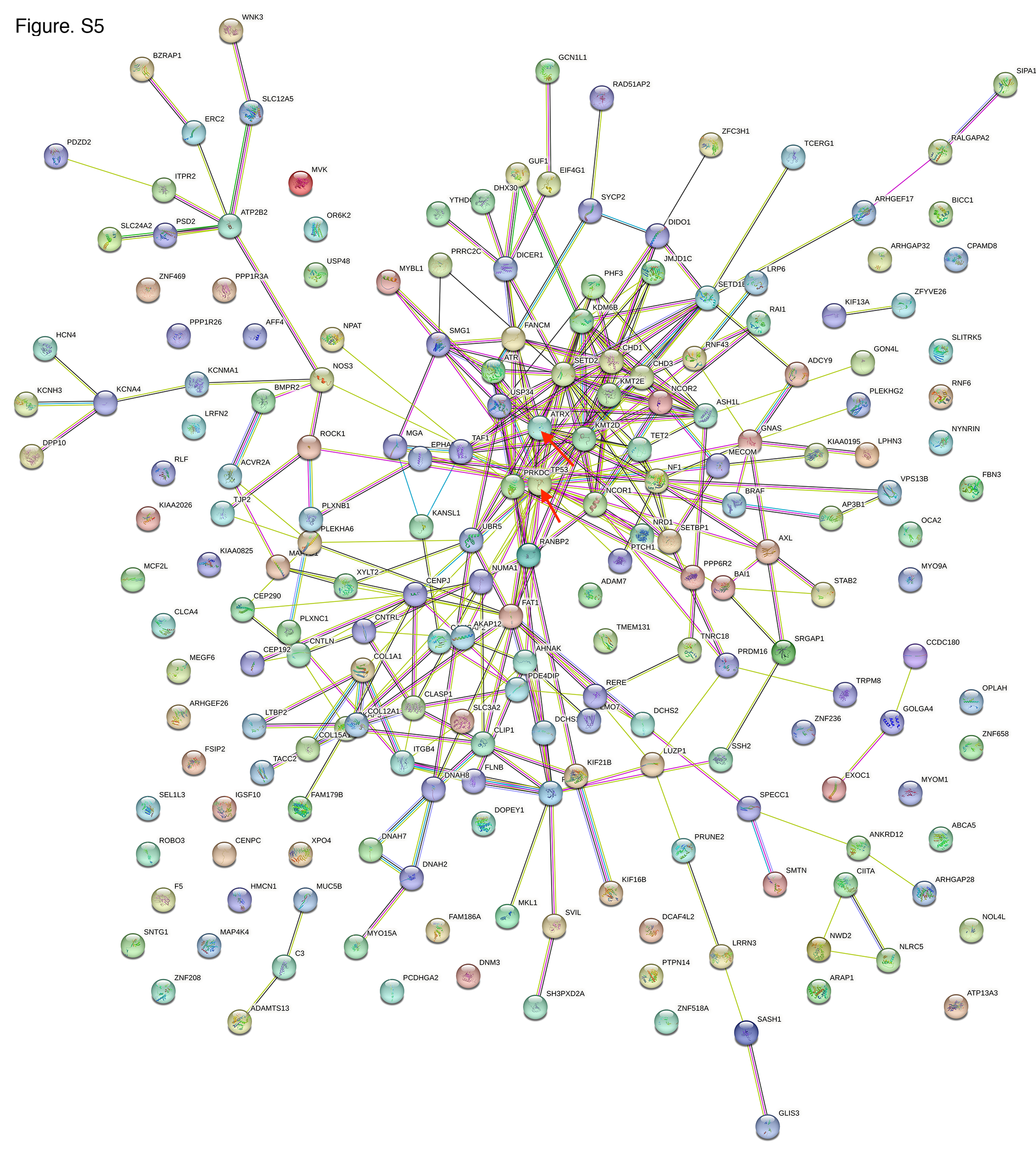

**Fig. S5, The PPI network of TP53, ATRX, and the top 200 potential regulators of XAR-PIX.** TP53 and ATRX were indicated by red arrows.

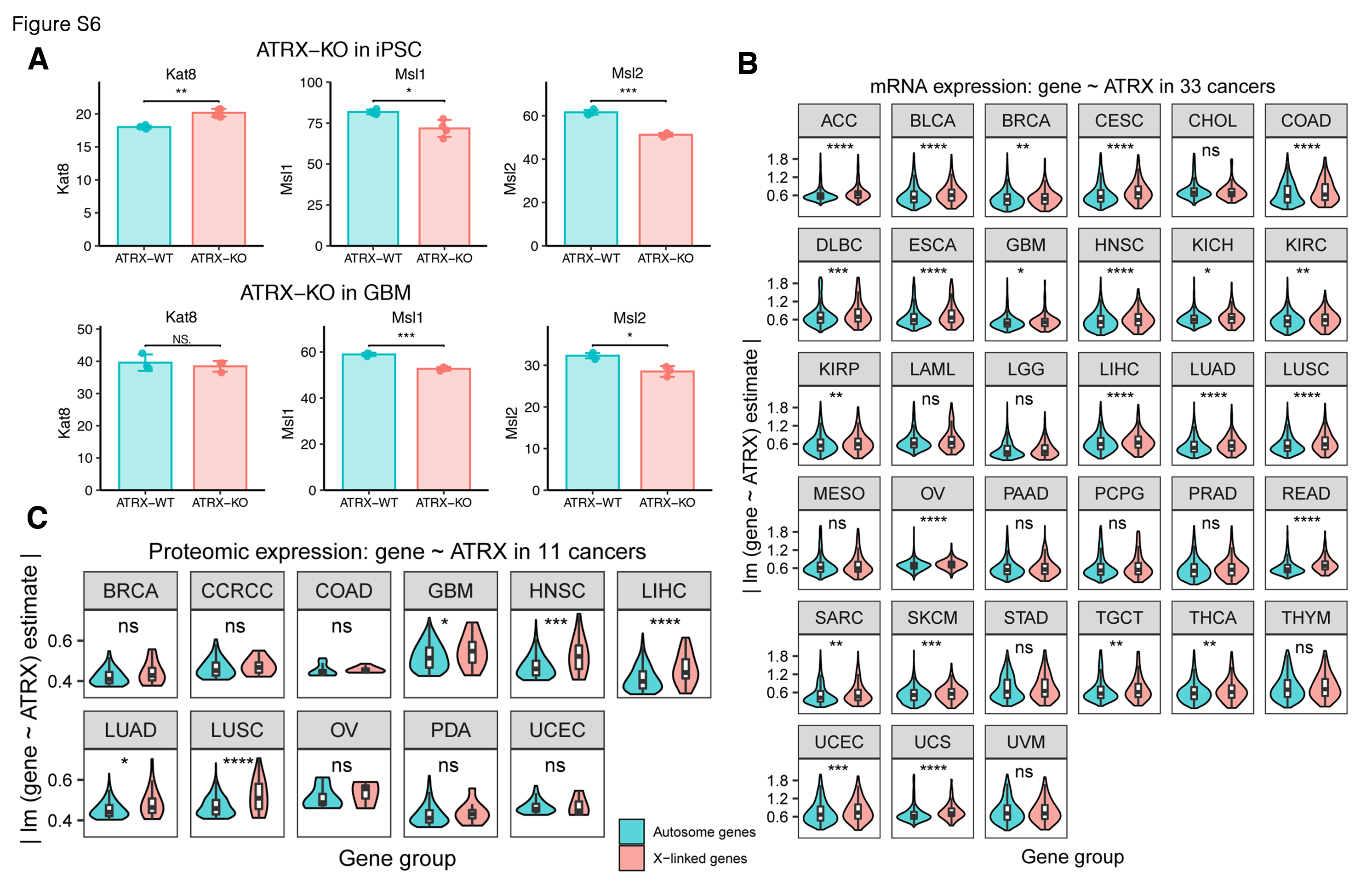

**Fig. S6, *ATRX* regulates XAR**. (**A**) Expression levels of KAT8, MSL1, and MSL2 after ATRX-KO in iPSC (top) and GBM (bottom) cells. (**B**-**C**) For autosomal and X-linked genes, linear regression models (each gene expression ~ *ATRX* expression) were fit to obtain regression coefficients of *ATRX* in each tumor, using mRNA (**B**) or proteomic (**C**) expression data. Absolute values of *ATRX* regression coefficients were compared between the two gene groups. Larger absolute values of *ATRX* coefficients indicate higher impact on gene expression.

Two-sided Wilcoxon test. * *P*<= 0.05, ** <=0.01, *** <= 0.001, **** <= 0.0001.

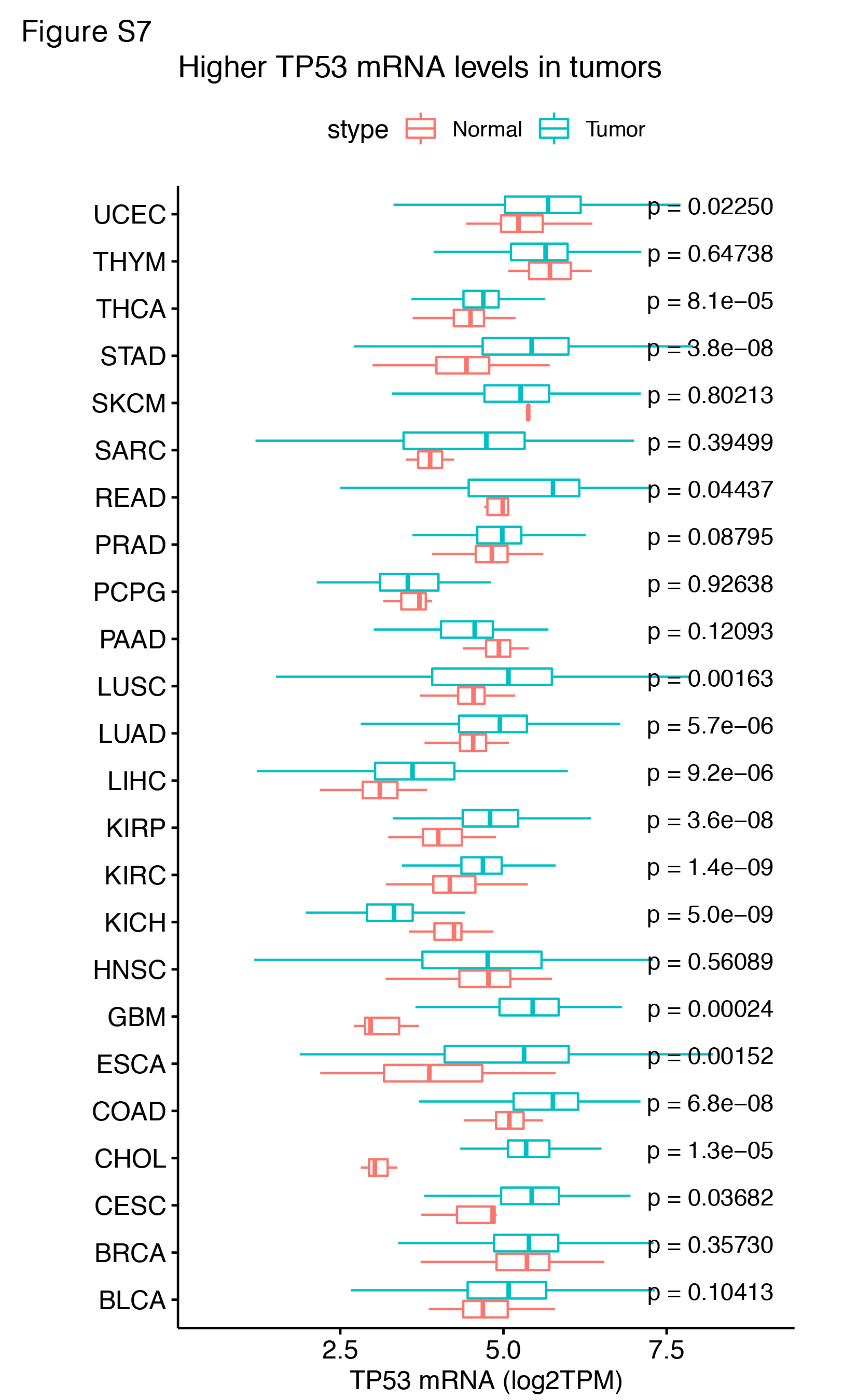

**Fig. S7, Comparison of TP53 mRNA levels across cancer types of TCGA.** Two-sided Wilcoxon test.

**Supplementary Table Legends**

**Table S1.** Gene classifications for X chromosome and autosomal genes.

**Table S2.** Enriched RNA-protein interactions for XAR-associated lncRNAs.

**References**

Loda A, Collombet S, Heard E. 2022. Gene regulation in time and space during X-chromosome inactivation. Nat Rev Mol Cell Biol 23:231-249.

Statello L, Guo CJ, Chen LL, Huarte M. 2021. Gene regulation by long non-coding RNAs and its biological functions. Nat Rev Mol Cell Biol 22:96-118.
